# Pannexin-1 regulates cellular identity and excitability in the developing cortex

**DOI:** 10.64898/2026.01.09.697058

**Authors:** Norma K. Hylton, David J. Kang, Steven C. Decker, David Exposito-Alonso, Christie N. Cambridge, Sean R. Golinski, Karla I. Soriano, Madeline C. Burke, Ximena Gomez-Maqueo, Qiurui Chen, Ruilong Hu, Jennifer E. Neil, Maya Talukdar, Rebecca E. Andersen, Trenton M. Buckner, Xuyu Qian, Anusha D. Doddi, Stephen R. Braddock, Chunhui Cai, Liang Sun, Ellen M. DeGennaro, Shyam K. Akula, Sergi Simo, Christopher A. Walsh, Richard S. Smith

**Affiliations:** Division of Genetics and Genomics, Boston Children’s Hospital, Boston, MA 02115; Departments of Pediatrics and Neurology, Harvard Medical School, Boston, MA, 02115; Harvard-Massachusetts Institute of Technology MD/PhD Program, Harvard Medical School, Boston, MA 02115; Howard Hughes Medical Institute, Boston Children’s Hospital, Boston, MA 02115; Department of Cell Biology and Human Anatomy, University of California Davis, Davis, CA, 95616; Allen Discovery Center for Human Brain Evolution, Boston Children’s Hospital and Harvard Medical School, Boston, MA, 02115; Department of Pharmacology, Feinberg School of Medicine, Northwestern University, Chicago, IL 60611; Broad Institute of MIT and Harvard, Cambridge, MA, 02142, USA; Division of Neurology, Department of Pediatrics, Children’s Hospital of Philadelphia, Perelman School of Medicine, University of Pennsylvania, Philadelphia, PA 19104; Division of Medical Genetics, Department of Pediatrics, Saint Louis University School of Medicine, St Louis, Missouri 63104; Research Informatics, Boston Children’s Hospital, Boston, MA, 02115; Division of Health Sciences and Technology, Massachusetts Institute of Technology, Cambridge, MA 02142, USA

**Keywords:** Cortical development, polymicrogyria, channelopathy, Pannexin-1, cell fate

## Abstract

Mutations in genes encoding a range of ion-conducting proteins disrupt development of the cerebral cortex in humans, often causing polymicrogyria (PMG); yet how ion conduction guides the development of cortical architecture is not clear. Here, we describe three individuals with brain malformations including PMG and microcephaly in whom *de novo,* missense mutations were identified in *PANX1*–encoding an ATP and ion conducting channel. We show that these PMG-associated PANX1 mutations (p.D14H, p.M37R, and p.N338T) disrupt normal glycosylation and confer gain-of-function with respect to ATP release and channel conductance. *In vivo* modeling of mutant PANX1 in cortical progenitor cells demonstrated disrupted cell migration and excess cell death in both mice and ferret models. *In vitro* modeling of the p.N338T allele in human induced pluripotent stem cell (hiPSC)-derived neurons further revealed changes in neuronal morphology and increased excitability, and dysregulation of key transcription factors critical for neurogenesis and cell fate. In addition, p.N338T iPSC-derived neural progenitor cells displayed increased PANX1 activation. Our results show that normal PANX1 function contributes to cortical structure through regulation of ion conductance and ATP release and provides insight into how these processes influence corticogenesis and cytoarchitecture more broadly.

**Significance Statement:** The purpose of ionic flux in human cortical histogenesis remains poorly examined. The recent identification of mutations in the gene encoding the ATP-releasing ion channel Pannexin 1 (PANX1) associated with polymicrogyria (PMG) provides a window for investigating processes of brain development regulated by ion and metabolite flux. In this study, we apply a combination of *in vitro* and *in vivo* approaches to both establish the pathogenicity of *PANX1*-associated PMG mutations and elaborate how perturbations in PANX1 disrupt cortical architecture. Our findings not only provide insight into the pathogenesis of PMG, but also further establish that conductive membrane properties of cortical progenitors influence neuronal identity and migration during neurogenesis.

## Introduction

Mutations in ion channel genes controlling action potential generation in mature neurons contribute to a large fraction of pediatric and adult epilepsy by shifting electrical activity (1–3), but it has been more recently appreciated that mutations in other ion channel-encoding genes with early embryonic expression also cause cortical malformations. These mutational effects demonstrate that ion channels do not merely influence the function of mature neurons after development, but also have additional, still poorly understood, functions that regulate the developing brain (4). Electrical activity and ionic fluxes have been reported at early stages of cortical development (5, 6), often in the absence of action potentials, and have been shown to influence neural progenitor cell proliferation (7–9), differentiation (10), and migration (11, 12).

Ionic flux and excitability are essential regulators of neurogenesis. Membrane potential oscillations and depolarization events regulate cell cycle progression (6, 7, 9, 13–17), while the sequential generation of neuronal cell types in the cortex from apical progenitors has been linked to progressive hyperpolarization of these progenitor cells (18). During prenatal development, before synapse formation and action potential propagation, progenitor cells exhibit slow depolarizing calcium (Ca^2+^) transients (5, 6, 19), and transient synapse formation between immature neurons has been shown to guide migration along radial glia (20, 21). Neuronal cell identity also hinges on spontaneous developmental activity (22–24), where ion channel expression and excitability promote the specification of cell types, including the differentiation of hippocampal progenitors (8), and the fate of cortical interneurons (12).

Disruptions of neural progenitor ionic flux and excitability influence corticogenesis, as demonstrated by cortical malformations caused by mutations in genes encoding ion-conducting proteins (4, 25). Recently, we reported whole exome sequencing (WES) results from individuals with polymicrogyria (PMG) and found that nearly 20% of variant-containing genes in solved cases encoded ion-conducting proteins (25). While studies of ion flux in cortical development have largely focused on voltage- or ligand-gated channels with well-defined synaptic and action potential functions (26–28), other channels expressed in developmental cell types also influence cellular excitability and cortical development. One such channel is Pannexin-1 (PANX1), a large-pore homo-heptameric plasma membrane glycoprotein broadly expressed in neurons and glia (29–31) that supports the flux of ATP and other large signaling metabolites (32). PANX1 regulates progenitor proliferation (30, 33), migration (34), and survival (35), yet its contribution to *in vivo* cortical histogenesis and cortical malformations remains unclear. Moreover, although purinergic signaling influences cell proliferation, including in intermediate neural progenitors (36–40), the contribution of PANX1 to ATP release in neural progenitors has not been examined.

Here we describe three individuals with *de novo* missense mutations in *PANX1* who presented with polymicrogyria (25), confirming a role for normal PANX1 activity *in utero* in cortical development. Inherited, activating variants in *PANX1* have been previously associated with female infertility (41–45), but without evidence of abnormal cortical development. In this study, we demonstrate PMG-associated mutations cluster in the channel pore, are predicted to disrupt gating, alter glycosylation, and confer gain-of-function activity through increased ionic flux and ATP release *in vitro*. In mice and ferrets, *in utero* electroporation of mutant PANX1 causes cell-autonomous and non-cell-autonomous neuronal migration defects and cell death. Proband iPSC-derived neurons exhibit increased firing and coincident discharges, accompanied by dysregulation of transcription factors critical for cell identity and signaling, while proband-derived neural progenitor cells (NPCs) demonstrate increased channel permeation, further supporting pathogenic gain-of-function. Our results demonstrate how disrupted PANX1 function may alter cortical structure and emphasize the importance of tightly regulating ionic flux and cellular excitability in progenitors and migrating neurons.

## Results

### Individuals with predicted pathogenic *PANX1* mutations have severe cortical malformations

In a previously reported WES brain malformation cohort study (25), we identified three unrelated individuals with severe PMG and *de novo* missense mutations in Pannexin-1 (*PANX1*; NM_015368.4) (Table S1). Individual PMG20601 [female, *PANX1* c.40G>C, p.(Asp14His)] presented with congenital microcephaly, infantile spasms, partial myoclonic seizures, cortical visual impairment, and global developmental delay. Brain MRIs at 12 and 26 months demonstrated extensive bilateral PMG centered in the perisylvian region and extending over the convexities, severe callosal agenesis, markedly depressed white matter, decreased thalamic volume, mild inferior vermian hypoplasia, and thinning of the brainstem (Figure 1A). PMGSL101 [female, *PANX1* c.110T>G, p.(Met37Arg)] exhibited mild acquired microcephaly, spastic quadriplegia, bilateral hyperopia, and global developmental delay. There were no documented seizures as of 6.5 years of age. Brain MRI at 5 months revealed extensive bilateral perisylvian PMG extending over the frontal and temporal lobes, moderately depleted white matter volume and reduced thalamic volume (Figure 1B). PMG24901 [male, *PANX1* c.1013A>C, p.(Asn338Thr)] presented with acquired microcephaly, right esotropia and myopia, left hemiparesis, and global developmental delay. Seizures developed around 3 years of age. Brain MRI at 10 months showed a small right hemisphere with diffuse PMG, reduced white matter and thalamic volume, and prominent perivascular spaces (Figure 1C). The presence of a left-sided cortical malformation was difficult to appreciate.

**Figure 1:**
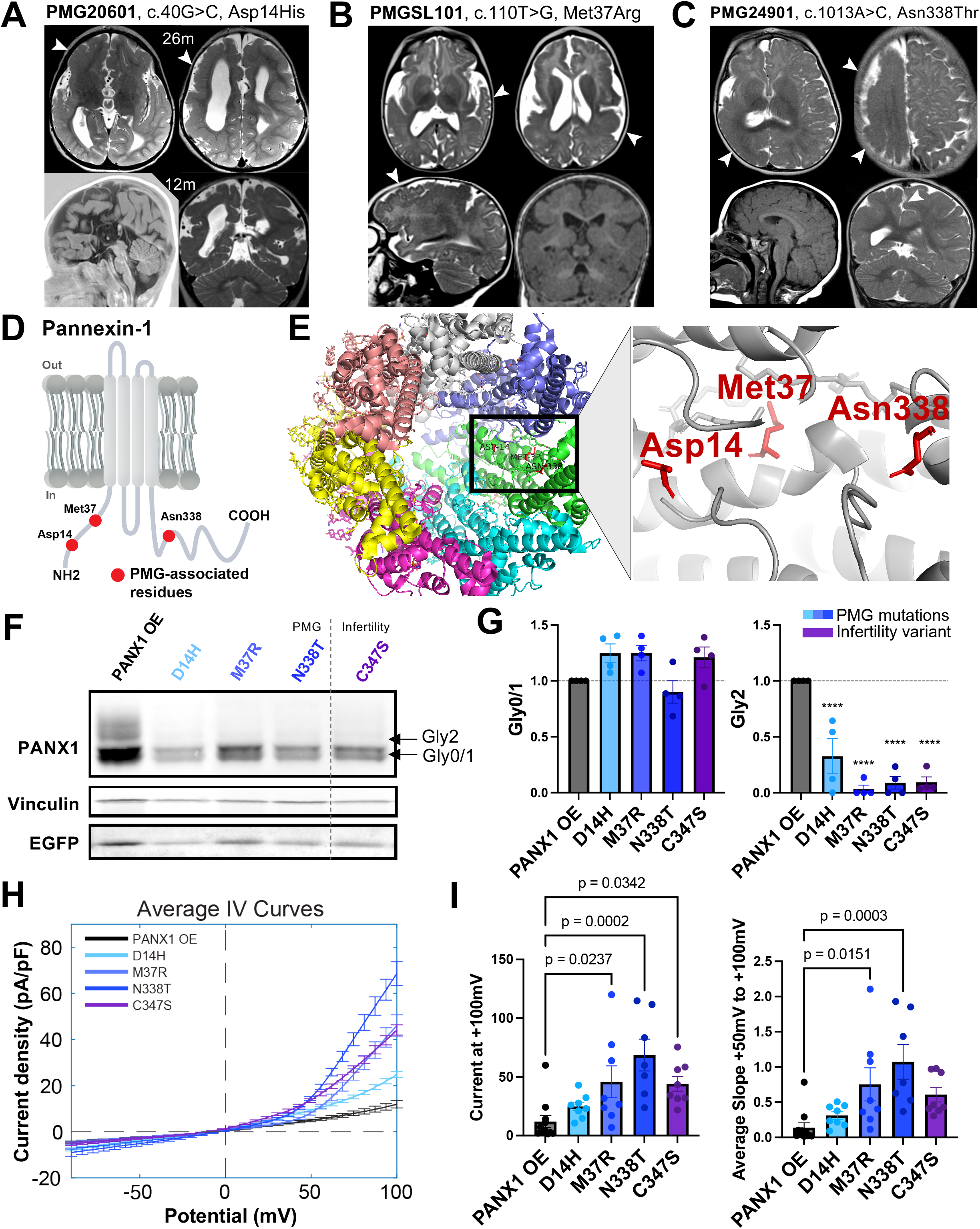
*De novo* mutations in *PANX1* identified in individuals with polymicrogyria and microcephaly disrupt complex glycosylation and increase ionic conductance. **(A-C)** Representative MRIs of affected individuals reveal cortical malformations, PMG, and abnormal gyral folding. **(A)** PMG20601, 12 months (m) and 26m: extensive bilateral PMG, centered in the perisylvian region and extending convexities, severe callosal agenesis, markedly reduced white matter, decreased thalamic volume, mild inferior vermian hypoplasia, and thinning of the brainstem. Bottom left image obtained from T1W_IR MRI series. **(B)** PMGSLU101, 5m: extensive bilateral perisylvian PMG extending over the frontal and temporal lobes, moderately depleted white matter volume and reduced thalamic volume. **(C)** PMG24901, 10m: small right hemisphere with diffuse PMG, reduced white matter and thalamic volume, and prominent perivascular spaces. See Table S1. **(D)** Schematic of PANX1 with red colored amino acids indicating locations of each mutation. See Figures S1 and S2. **(E)** *Right*, heptameric cryo-EM structure of PANX1 from Ref. (48) rendered in PyMol (6WBF). Three PMG-associated amino acid residues, Asp14, Met37, and Asn338 (red), are in indicated in the same PANX1 subunit of the heptamer. *Left*, close up of the domain where amino acids reside. **(F)** Western blotting for PANX1 (∼55kDa) reveals three known glycosylated species: Gly0 (unglycosylated), Gly1 (mannose), and Gly2 (complex glycosylation). All three PMG mutations significantly reduce Gly2. **(G)** Quantification of 4 western blots from 4 independent transfections (ANOVA with post-hoc Tukey test, **** p< 0.0001). PMG mutations in blues, infertility in purple. All values normalized to transfection level (EGFP) and membrane protein loading control (Vinculin). No significant difference in Gly0/1 expression. **(H)** Average I-V (current voltage) relationship curves of HEK293Ts expressing either WT *PANX1* (OE) or MT *PANX1* from whole-cell patch clamping using a ramp protocol (-90mV to +100mV, see Figure S9). M37R, N338T, and C347S demonstrate increased conductance at depolarized potentials relative to WT channels (n = 8 for each condition). **(I)** *Left*: current density at +100mV is significantly increased for M37R, N338T, and C347S. *Right*: the average slope of the curve from +50mV to +100mV is significantly increased for M37R and N338T (ANOVA with post-hoc Tukey test). Mean +/- SEM in **(H)-(I)** presented.

Protein prediction tools suggest that all three missense mutations alter PANX1 activity. D14 and M37 are located near the N terminus and N338 near the C terminus (Figure 1D; Figure S1, S2), regions with ascribed roles in channel gating and ion permeation, where electrostatic changes introduced by these mutations may alter gating mechanisms (46–50). AlphaMissense, which incorporates AlphaFold structural information (51), predicts all three mutations to be pathogenic, with scores of 0.921 (D14H), 0.9011 (M37R), and 0.9448 (N338T), corroborated by additional *in silico* tools (Table S1). The affected amino acids are highly conserved, particularly among mammals (Figure S1), and cluster within the channel pore in a likely-open cryo-EM structure of PANX1 (48) (Figure 1E), supporting a shared mechanism of perturbed channel function. Together, the similar MRI phenotypes, pore clustering, *de novo* nature, and electrostatic changes strongly suggest that these *PANX1* cause channel dysfunction and cerebral cortical malformation.

### *PANX1* expression is enriched in the prenatal human cortex

Both *PANX1* and *PANX2* are expressed in the human brain (30), yet only *PANX1* demonstrates temporal enrichment during gestational development in bulk RNA sequencing of human cortex from the Allen Developing Human Brain Atlas (Figure S3A). RNAscope at gestational week (GWs) 19 shows high PANX1 expression in the cortical plate and ventricular zones (Figure S3B), while MERFISH at GWs 18, 20, and 21 demonstrates a similar pattern, with a slight increase coinciding with late neurogenesis and formation of layer 2/3 neurons (52) (Figure S3C-D). Single-cell RNA-sequencing from a human cortical development atlas (53) similarly demonstrates fetal enrichment of *PANX1* across progenitor subtypes, neurons, and glia in the prefrontal cortex and area V1 (Figure S4A-D), consistent with previous work (30, 54); *PANX1* is also broadly expressed in ferret cortex, with some enrichment in deep-layer neurons (Figure S5). Although previous work detected PANX1 immunoreactivity in the CP and SVZ, but not VZ, at mid-gestation (30), we find that PANX1 immunoreactivity declines prenatally, we find higher expression at GW 15 than GWs 20 and 34, particularly in the VZ and SVZ (Figure S6A-C), a pattern recapitulated in protein from human fetal synaptosomes (Figure S7). Together, these results indicate that PANX1 is preferentially expressed during corticogenesis across multiple cell types, including cortical progenitors and migrating and differentiating neurons.

### *PANX1* mutations disrupt complex glycosylation and increase ATP release

We first investigated the identified *PANX1* mutations *in vitro* to determine their effects on protein glycosylation, as infertility-associated *PANX1* mutations have been shown to disrupt glycosylation (42–45). We generated piggyBac constructs expressing human *PANX1* and *EGFP* linked by a T2A peptide (55), which enable expression following transposon integration using a hyperactive transposase (piggyBase) (56) (Figure S8A). We also generated a construct containing the previously reported infertility-associated gain-of-function variant C347S (Table S2 and Figure S1B) (41), which reduces the heavily glycosylated Gly2 form of PANX1, a feature associated with gain-of-function; notably, residue N338 has also been predicted to be a PANX1 glycosylation site (41). Western blotting of transfected HEK293Ts demonstrated that all three PMG-associated mutations significantly reduced or eliminated Gly2 expression (Figure 1F-G), indicating reduced complex glycosylation (Figure S8B-C).

Given that previous glycosylation-disrupting variants were also shown to be gain-of-function (42–45), we next assessed whether our newly identified PMG-associated mutations might display disrupted hemichannel activity. PANX1 forms hemichannels that release ATP extracellularly (57), and we tested whether PMG mutations alter ATP release by transfecting mouse neuroblastoma (N2A) cells with either WT or mutant (MT) PANX1 and collecting the extracellular media. PMG mutations significantly increased the release of ATP, and this effect was attenuated by pre-treatment with the hemichannel blocker carbenoxolone (CBX) (Figure S8D).

We next assessed PANX1 MT channel conductance by whole-cell patch clamp using voltage ramps from −90 to +100 mV in *PANX1*-transfected HEK293T cells. M37R, N338T, and C347S exhibited increased ionic conductance and higher maximum current densities at depolarized potentials compared with WT *PANX1*, whereas D14H was not significantly different (p = 0.67) (Figure 1H-I and Figure S9). M37R and N338T also showed greater sensitivity to membrane potential, with significantly steeper slopes between +50 and +100 mV, indicating increased ionic flow and a more rapid increase in current density with depolarization.

### PANX1 overexpression *in vivo* impedes mouse neuronal migration and induces cell death

To examine the effects of *PANX1* gain-of-function on neocortical migration and cell fate, we performed in utero electroporation (IUE) at E14.5 to express WT-PANX1-EGFP, N338T-PANX1-EGFP, or EGFP control, co-electroporated with an episomal dsRed plasmid (Figure 2A). Two days later, fresh tissue sections underwent 24-hour time-lapse confocal imaging to track EGFP+ cells migrating toward the CP (Figure 2B and Video S1-3). We quantified migration speed and path straightness, defined as the ratio of displacement to distance traveled, and found that EGFP-expressing cells in N338T-IUE brains migrated significantly slower and followed altered trajectories compared with both EGFP control and WT-PANX1 cells (Figure 2C). These results suggest that PMG-associated PANX1 mutations result in neuronal migration defects in the developing cortex.

**Figure 2:**
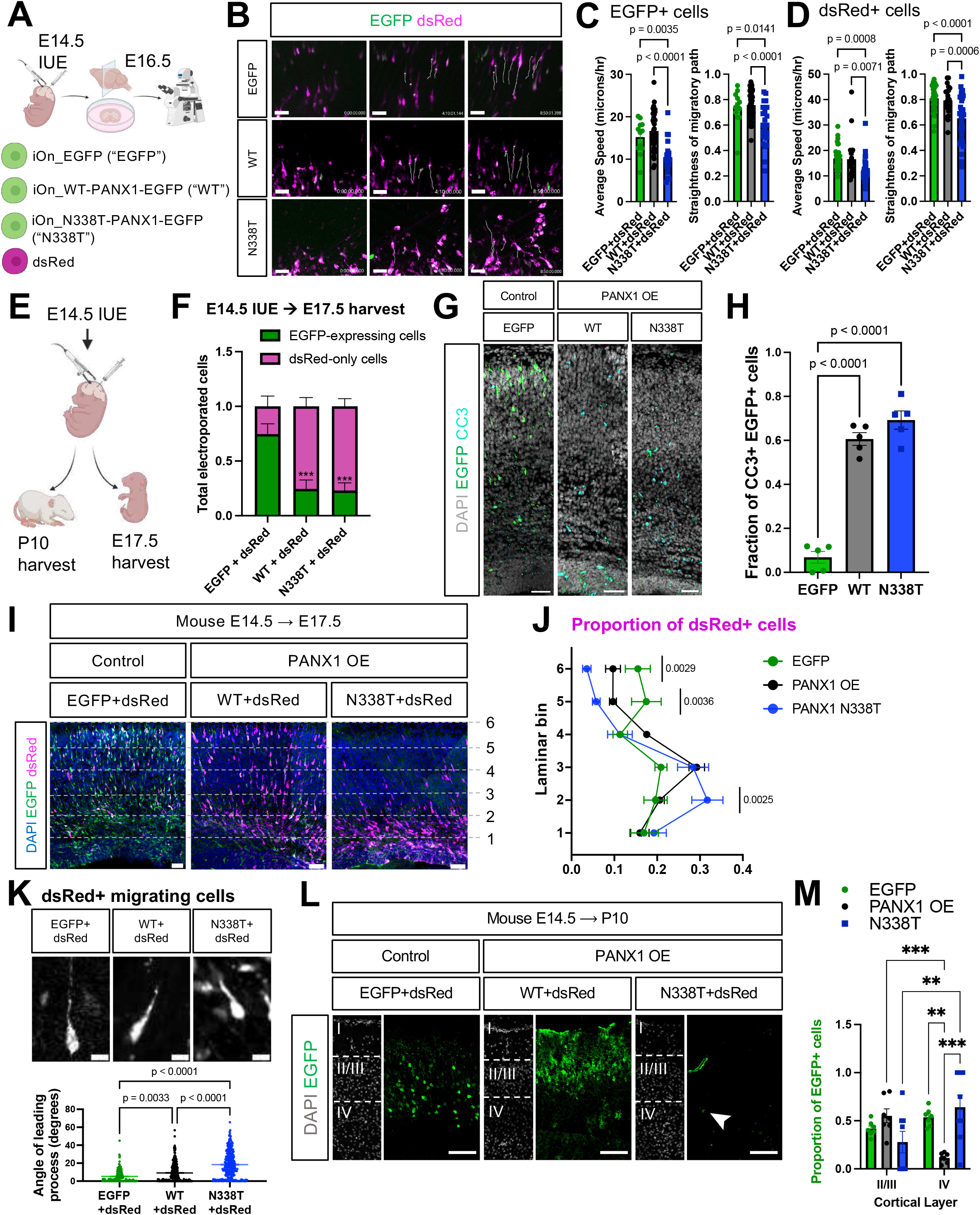
Overexpression of mutant *PANX1* disrupts cell-autonomous and non-cell-autonomous neuronal migration and causes cell death. **(A)** Schematic of E14.5 mouse co-electroporation of integrating *PANX1*-EGFP constructs with episomal pCAG-dsRed for time-lapsed confocal imaging for 24-hrs. **(B)** Representative images from videos of migrating electroporated cells. Scale bar = 30μm. **(C)** Average speed (μm/hr) and straightness of migratory paths across conditions for EGFP+ cells. EGFP+ cells expressing N338T *PANX1* demonstrated reduced speed and deviated migration (ANOVA with post-hoc Tukey) compared to EGFP and WT. **(D)** Average speed (μm/hr) and straightness of the migratory path for dsRed-only expressing cells. Results indicate reduced migration speed and deviations in migratory path of dsRed-expressing cells in N338T-electroporated mice (ANOVA with post-hoc Tukey). Dot = individual cell; data obtained from >20 cells per condition in at least 3 mice. **(E)** Schematic of mouse E14.5 IUE, harvested at either E17.5 (n = 5 EGFP, n = 8 WT, n = 8 MT) or P10 (n = 5 EGFP, n = 7 PANX1 WT, n = 9 N338T). **(F)** *PANX1*-electroporated mice demonstrate significant reductions in EGFP-expressing cells compared to EGFP control + dsRed, p = 0.0008 WT, and p = 0.0004 N338T, two-way ANOVA with Sidak multiple comparisons test. **(G, H)** In E17.5 electroporated mice, there is a significant increase in the number of Cleaved caspase-3 (CC3+) electroporated cells in both WT and N338T electroporated conditions compared to EGFP+dsRed (n = 5 EGFP, n = 5 PANX1 WT, n =6 N338T; scale bar = 50μm). ANOVA with post-hoc Tukey test. OE, overexpression. **(I, J)** Mice electroporated with mutant *PANX1* demonstrated dsRed+ cells with a laminar positioning defect, with more cells concentrated in the VZ/SVZ compared to EGFP and WT conditions at E17.5, scale bar = 50μm. Cells counted and binned according to migratory position by dividing cortex into 6 equally sized laminar bins. Two-way ANOVA with post Sidak multiple comparisons tests revealed more electroporated cells found in bin 2 (p = 0.0025), and fewer cells in bins 5 (p = 0.0036) and 6 (p = 0.0029) compared to EGFP alone. **(K)** *Top*, the angle of the leading process of migrating dsRed+ cells was measured from vertical. *Bottom*, WT PANX1 and N338T conditions demonstrate significant increases in angle deviation (p = 0.0033 EGFP vs. WT; p < 0.0001 EGFP vs. N338T; p < 0.0001 WT vs. N338T, one way ANOVA with post hoc Tukey test). >100 cells counted per condition. Scale bar = 10μm. **(L, M)** P10 mouse neurons targeted by EGFP control migrate to layers 2/3 and 4. WT *PANX1* demonstrate layer 2/3 predominance (p = 0.0008, two-way ANOVA with post-hoc Tukey test), with a significant reduction in layer 4 neurons compared to EGFP control (p = 0.0031) and N338T (p = 0.0002). EGFP+ neurons targeted by N338T *PANX1* are shifted to layer 4 of the cortex compared to layer 2/3 (p = 0.0026). Scale bar = 100μm.

To assess non-cell-autonomous effects, we also tracked dsRed+ cells lacking *PANX1* overexpression and found significantly reduced migration speed and altered trajectories in N338T-electroporated brains (Figure 2D), suggesting that mutant PANX1 also disrupts migration within the environmental niche. During time-lapse imaging, cells expressing WT or N338T *PANX1* also exhibited apparent cell death and disorganized, non-laminar activity (Video S2-3), prompting us to assess cell loss and cytotoxicity following IUE at E14.5 and subsequently harvesting tissue at either E17.5 or postnatal day 10 (P10) (Figure 2E). At E17.5, the fraction of EGFP+ cells was significantly reduced in both WT and N338T conditions compared with EGFP control (Figure 2F), and cleaved caspase-3 (CC3) staining confirmed increased apoptosis (Figure 2G, H). While few control EGFP+ cells were CC3+, approximately 60% and 70% of WT- and N338T-expressing cells, respectively, were CC3+ (Figure 2H), demonstrating that overexpression of either WT or MT *PANX1* increases apoptosis in the developing mouse cortex.

Given the degree of EGFP+ cell death, we examined whether the positioning of dsRed+ cells lacking PANX1 expression was also disrupted. At E17.5 following E14.5 IUE, N338T-*PANX1* electroporation resulted in significantly fewer dsRed+ neurons reaching the cortical plate and altered laminar positioning compared with EGFP control (Figure 2I, J, and Figure S10A). WT-*PANX1* produced an intermediate phenotype, with nonsignificant trends toward reduced dsRed+ cells in laminar bins 5 (p = 0.074) and 6 (p = 0.23). We next measured the angle of leading processes of dsRed+ migrating cells and identified significantly deviated angles from vertical following either WT or N338T *PANX1* electroporation, with greater deviations in N338T brains (Figure 2K). Together, these findings support non-cell-autonomous disruptions to cell migration following perturbation of PANX1 function in the developing mouse cortex.

To determine whether these early positioning changes persist after migration is complete, we analyzed brains at P10 following E14.5 IUE (Figure 2E and Figure S10B). N338T-electroporated brains again showed significantly fewer EGFP+ cells than controls (Figure S10C), consistent with widespread cell death. Whereas control EGFP+ neurons were distributed approximately evenly across layers 2/3 and 4, WT-*PANX1* brains showed depletion of layer 4 neurons and enrichment in layers 2/3, while N338T IUE brains demonstrated a predominance of layer 4 EGFP+ neurons (Figure 2L-M). These altered distributions may reflect changes in neuronal fate or persistent mis-migration.

### Overexpression of mutant *PANX1* in ferrets recapitulates cell positioning deficits and cell death

To extend our mouse findings to a gyrencephalic mammal, we performed postnatal electroporation in ferrets using the same constructs as described above. P1 ferret visual cortices were electroporated under stereotactic guidance and harvested at P10 (Figure 3A). Gyrification of P10 brains was grossly indistinguishable between *PANX1* WT OE and N338T electroporated animals (Figure S11A), although gyrification in normal ferrets is rudimentary at this age (58). We first examined the proportions of PANX1-expressing electroporated cells and again noted a significant reduction in the fraction of pathogenic EGFP+ cells in N338T compared to WT OE (Figure 3B), suggestive of cell death (p<0.0037). Tissue staining with CC3 identified that 25-35% of cells overexpressing either WT or MT *PANX1* were positive for CC3+ with a nonsignificant trend towards an increase in the proportion of CC3+ electroporated cells from MT compared to WT (p = 0.1086, t-test; Figure S11B,C), likely due to cell death that has already occurred by P10 across both conditions.

**Figure 3:**
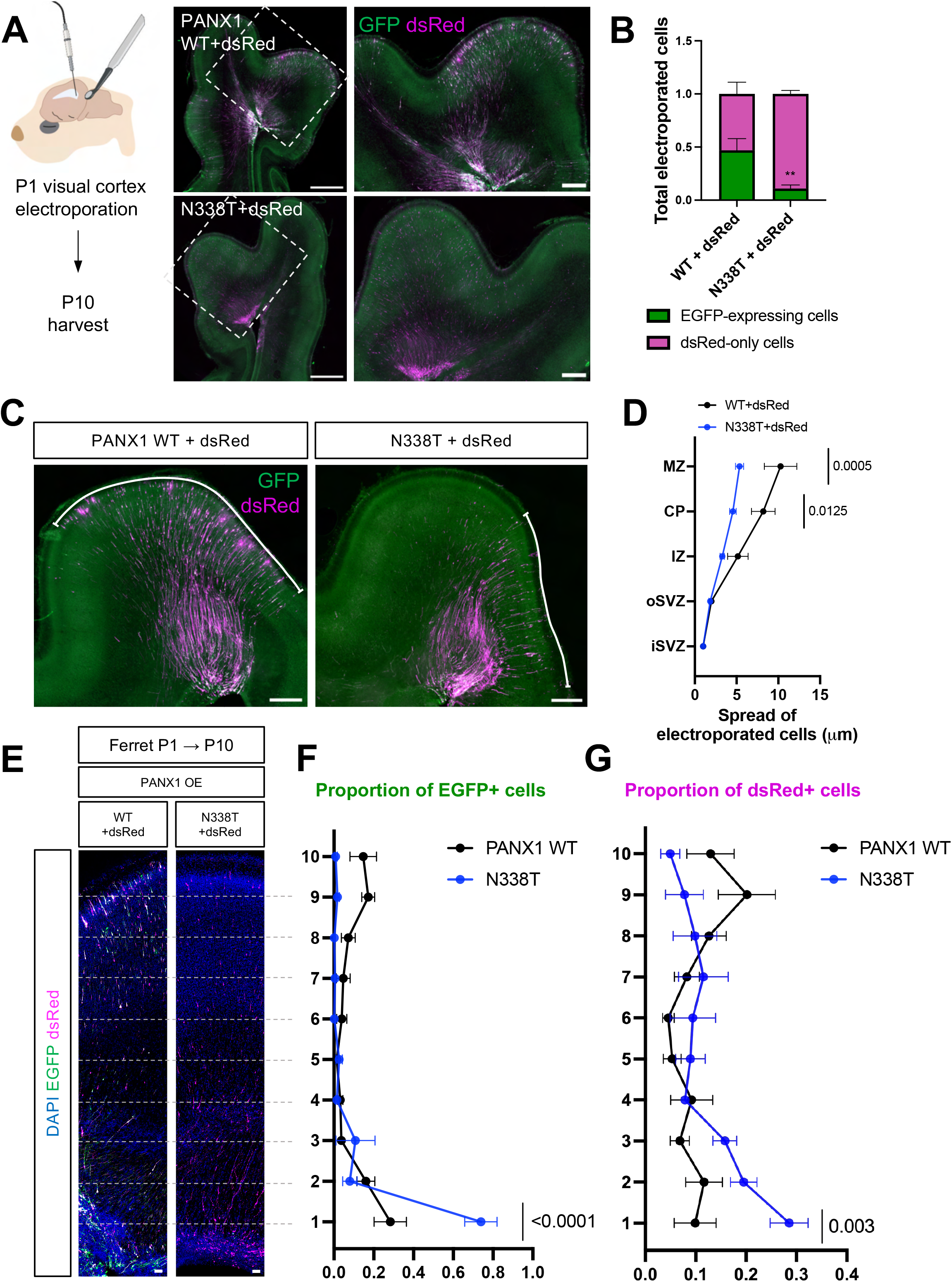
Overexpression of mutant *PANX1* in ferrets alters neuronal positioning and increases cell death. **(A)** *Left*, schematic of postnatal electroporation in visual cortex P1 ferret kits with plasmids as previously described (see Figure 2A). Ferrets harvested at P10 (n = 5 PANX1 OE, n = 5 N338T, for all panels), and >200 cells per brain analyzed. *Right*, representative confocal images of electroporated ferret visual cortex at P10 (left scale bar = 500μm; right scale bar = 200μm). **(B)** N338T-electroporated EGFP+ cells are significantly reduced in P10 ferrets (two-way ANOVA with Sidak multiple comparisons test, ** p = 0.0037). **(C, D)** WT-electroporated animals have greater tangential spread of electroporated cells compared to N338T (two-way ANOVA with Sidak multiple comparisons test). Spread determined by measuring curvilinear distance between electroporated cells within a gyrus. Scale bar = 200μm. **(E, F, G)** Electroporation of mutant *PANX1* in the developing ferret visual cortex results in disrupted neuronal migration, with more EGFP+ and dsRed+ cells present closer to VZ/SVZ (scale bar =200μm). Cells counted and binned according to migratory position by dividing cortex into 10 laminar bins. Two-way ANOVA with Sidak multiple comparisons tests revealed **(F)** more EGFP+ cells from N338T brains in laminar bin 1 (p < 0.0001) compared to WT and **(G)** more dsRed+ cells from N338T condition found in bin 1 (p = 0.003) compared to WT.

We next analyzed dispersion of migratory columns in *PANX1* WT- and N338T-electroporated ferrets. In gyrencephalic mammals, migrating neurons fan tangentially across developing gyri, a process associated with increased neurogenesis and mechanical forces that drive cortical folding (52, 59). Given the distorted cortical folding in our human PMG cases, we measured the curvilinear width (white line, Figure 3C) of migrating electroporated cells from the SVZ to MZ, normalized to the initial electroporated area (Figure 3C). WT-electroporated cells showed greater tangential spread across a gyrus than N338T cells, particularly in the CP and MZ (Figure 3D). This pattern suggests disrupted gyral expansion through reduced fanning of migrating neurons in N338T-electroporated ferrets.

We next quantified laminar positioning of EGFP+ and dsRed+ cells across 10 bins spanning the developing ferret visual cortex. Consistent with our mouse findings, N338T-electroporated brains showed significantly altered positioning of both EGFP+ and dsRed+ cells, with a greater proportion retained in the first laminar bin (Figure 3E-G and Figure S11D). Together, these findings demonstrate that mutant *PANX1* expression alters migration through both cell-autonomous and non-cell-autonomous effects on neuronal positioning, as well as reduced tangential spread in a gyrencephalic mammal.

Finally, we performed E32 IUE in the ferret somatosensory cortex using the same *PANX1* and dsRed constructs and harvested brains at P1 or P21. At P1, N338T-electroporated brains lacked gross EGFP expression (Figure S12), providing further evidence of PANX1-mediated cell death. In 15 N338T-electroporated ferret brains harvested at P21, none demonstrated gross EGFP or dsRed expression.

### Mutant iPSC-derived neurons have reduced neurite outgrowth and increased excitability

To assess neuronal phenotypes of gain-of-function PANX1 without overexpression, we generated NGN2-induced neurons from an iPSC line derived from PMG24901 (PMG24901-A; p.N338T) and two CRISPR-corrected isogenic controls (WT-T338N-1 and WT-T338N-2; Figure 4A). PMG24901 was the only individual out of the cohort of three who provided consent for iPSC generation, and derived iPSCs exhibited normal morphology and karyotype (46XY), pluripotency marker expression, and tri-lineage differentiation (Figure S13). Following NGN2-mediated differentiation into forebrain glutamatergic neurons (Figure 4A; see Methods), N338T neurons showed significantly reduced neurite outgrowth and branching compared to isogenic controls (Figure 4B). We next assessed cellular physiology using whole-cell patch clamp and multielectrode array (MEA) (Figure 4A, C-E). Minimal-stimulation current clamp recordings revealed similar action potential kinetics between WT and N338T-carrying neurons, including threshold potential and halfwidth duration (Figure 4C; p > 0.25 for each readout). Sodium influx, potassium efflux, capacitance, and membrane resistance were also comparable across conditions.

**Figure 4:**
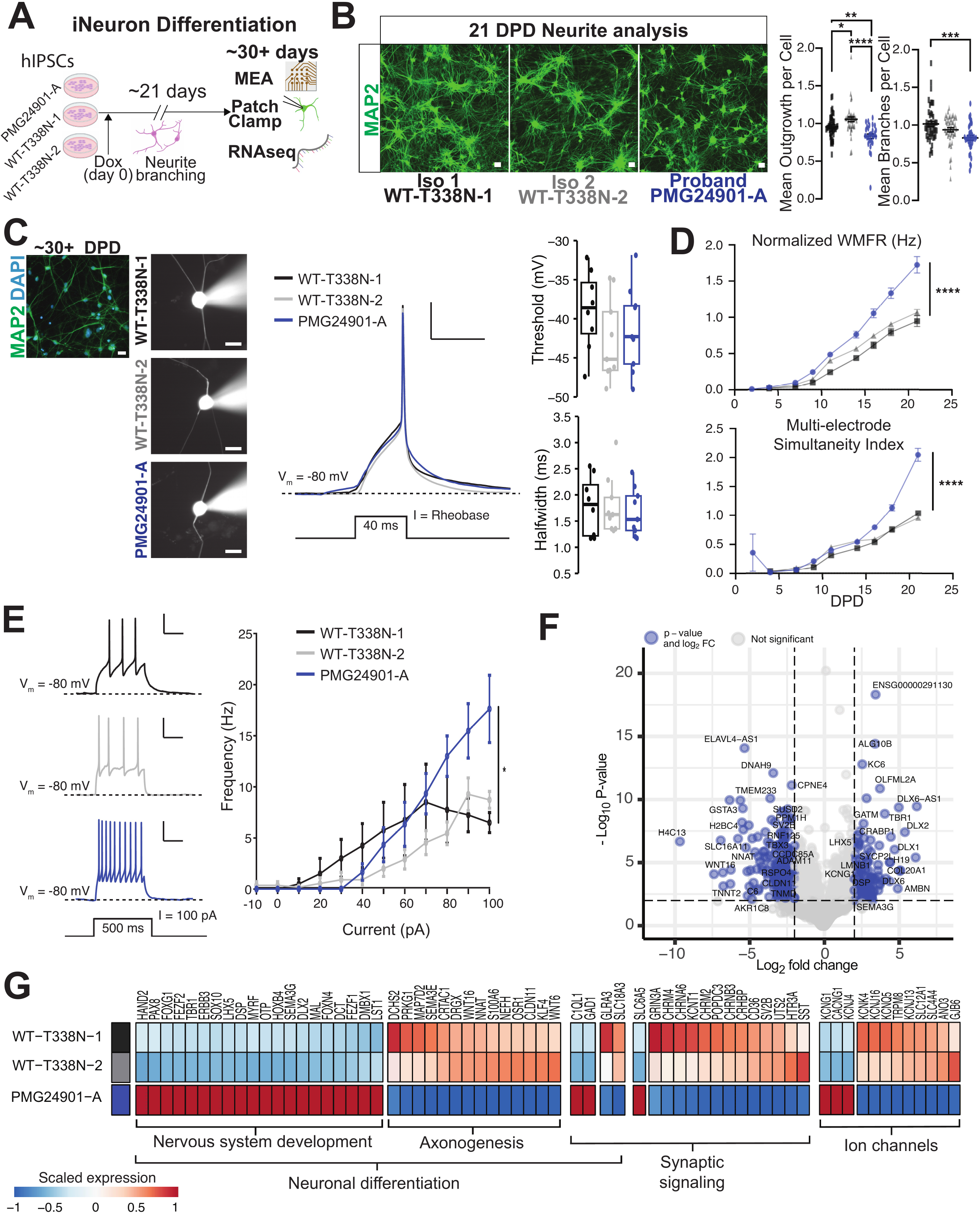
PMG-associated mutations alter neuronal morphology and excitability. **(A)** Schematic of NGN2 viral transduction of proband-iPSCs and two isogenic controls produced by CRISPR-mediated correction of PMG24901-A to generate homogenous cultures of induced neurons for downstream assays. **(B)** *Left*, representative images of MAP2+ NGN2-induced neurons from each cell line demonstrating reduced branching of PMG24901-A neurons. Scale bar = 400μm. *Middle*, automated analysis reveals a reduction in neurite length in PMG24901-A neurons compared to controls (** p = 0.0086, **** p < 0.0001, ANOVA with post-hoc Tukey test; >30 cells per condition). WT-N338T-1 with slightly reduced neurite length compared to WT-T338N-2 (p = 0.0371). *Left,* automated analysis of total branching further indicates a reduction in the overall number of branches in mutation-carrying neurons compared to WT-T338N-1 (p = 0.0004, ANOVA with post-hoc Tukey test; >30 cells per condition). No significant difference between WT-T338N-1 and WT-T338N-2 (p = 0.1913) or between WT-T338N-2 and PMG24901-A (p = 0.057). Data from three independent experiments; values normalized to an average of the two controls. **(C)** *Left*, representative dye-loaded neurons during patch experiments from each condition, scale bar = 25 μm. *Middle*, average single action potential of each condition generated by a 40 ms pulse at the rheobase current, scale bars = 20 mV, 50 ms. Neurons (n = 9 for PMG24901-A, n = 9 WT-T338N-1 and n = 8 for WT-T338N-2) were held near -80 mV with a holding current before being successively stimulated by pulses until rheobase value was reached. *Right*, threshold and halfwidth of neurons; no significant differences. Data presented as median +/- IQR with whiskers spanning the range of the data excluding outliers (two-sample, two-tailed Wilcoxon tests). **(D)** *Top*, normalized weighted mean firing rate reveals increased firing of mutation-carrying neurons compared to isogenic controls. *Bottom*, normalized multi-electrode simultaneity demonstrates significantly increased simultaneous discharges of mutant neurons. Data from three differentiations; 16 independent wells per cell line; values normalized to the averages of the maximum of the isogenic control. Curves fit to sigmoidal curve using a nonlinear regression model (**** p<0.0001, t-test of model parameters). **(E)** *Left*, representative neuronal spiking traces from each condition in response to a prolonged current injection of +100 pA for 500 ms; scale bars = 20 mV, 200 ms. *Right*, spiking frequency plotted as a function of input current from -10 pA to +100 pA in 10 pA steps. Proband neurons exhibited higher frequency firing at +100 pA compared to WT. Data as mean +/- SEM (* p = 0.0221 proband vs WT-T338N-1 and p = 0.0222 proband vs WT-T338N-2, two sample, two-tailed t-test), n = 7 for PMG24901-A, n = 8 for WT-T338N-1, and n = 13 for WT-T338N-2. **(F)** Volcano plot of differential expression analysis in DESEq2 depicting all tested genes. Blue circles are genes that passed both thresholds for significance (p < 0.01 and fold change |log2|> 2). Bulk RNA-seq differential gene expression performed on NGN2 neurons (n = 8, PMG24901-A; n = 6 WT-T338N-1; n = 8 WT-T338N-2). **(G)** Heatmap with scaled expression of select differentially expressed genes in PMG24901−A. Brackets highlight associated biological functions, detected through gene ontology enrichment analysis.

We next collected MEA recordings over 20–25 days post-differentiation (DPD) using independent neuronal differentiations co-cultured with human astrocytes, excluding wells with fewer than 10 active electrodes by 11 DPD (Figure S14A). We assessed weighted mean firing rate (WMFR) and multi-electrode simultaneity, a measure of the cross-correlative activity between electrode pairs, using nonlinear regression of data normalized to the average peak value. N338T neurons showed significantly increased peak WMFR and peak simultaneity compared with both WT lines, which did not differ from each other (Figure 4D). These findings provide robust evidence for augmented excitability, cellular firing, and simultaneous discharges due to *PANX1* mutation, consistent with seizures observed in PMG24901.

To further characterize cellular excitation, we generated current clamp excitability curves from minimal stimulation to maximal depolarization. N338T neurons fired higher-frequency action potential trains at high current stimulation (e.g., +100 pA), while firing frequencies were comparable to WT neurons at lower current injections (−10 to +90 pA; p > 0.05; Figure 4E). Acute treatment with the PANX1 antagonist mefloquine (1 µM) did not significantly dampen the excitability curve (Figure S14B), suggesting that increased excitation is not driven by acute PANX1 activation during stimulation.

We next performed bulk RNA-seq differential expression analysis of PMG24901-A NGN2 neurons and two isogenic controls using DESeq2 (60). While *PANX1* and *PANX2* expression were similar across genotypes (Figure S14C), 393 genes were differentially expressed in PMG24901-A neurons (Figure S14D,E). Among 146 upregulated genes, functional enrichment identified genes with CNS and embryonic development, regionalization, and patterning functions (Figure 4F; Figure S14F; Supplementary Dataset 2), driven in part by a high proportion of transcription factors (19%) involved in neurogenesis and cell fate, including *FOXG1*, *TBR1*, *FEZF1/2*, and *DLX1/2/5/6*. The 247 downregulated genes were enriched for neuronal processes including axonogenesis and synaptic signaling (Figure 4G), and included potassium channels (*KCNJ13, KCNJ16, KCNK4, KCNQ5,* and *KCNT1*) and core nucleosomal histones (*H2AC14, H2BC12L, H2BC13, H2BC17, H2BC18, H2BC4, H3C6, H4C13,* and *H4C8*; Figure S14F), with enrichment in senescence-related stress responses and *CHD1* transcriptional regulation (Supplementary Dataset 2). Together, these results demonstrate that *PANX1* mutation broadly alters gene expression networks governing developmental patterning, neurogenesis, and neuronal differentiation, and suggest that increased excitability may arise from shifts in ion channel expression rather than PANX1 channel activity alone.

### Patient-Derived NPCs Exhibit PANX1 Functional Hyperactivation

Considering *PANX1* expression in the human fetal SVZ and progenitor phenotypes following mouse and ferret IUE, we assessed PANX1 channel permeation in NPCs derived from PMG24901-A and CRISPR-corrected isogenic control hiPSCs (Figure 5A). NPC identity and PANX1 expression were confirmed by PAX6 and PANX1 immunostaining (Figure 5B and Figure S15A,B), with comparable PANX1 protein levels across genotypes by western blot (Figure 5C). We next measured PANX1 channel permeation using YO-PRO dye uptake under vehicle conditions or following stimulation with the purinergic receptor agonist BzATP (100 µM; Figure 5D). While baseline YO-PRO uptake was similar between proband and WT NPCs (p = 0.37 vs WT-T338N-1, p = 0.43 vs WT-T338N-2), BzATP induced significantly greater uptake in mutation-carrying NPCs (p < 0.01), with no difference between WT controls (p = 0.97; Figure 5E). Thus, despite comparable PANX1 protein expression, mutant NPCs exhibit increased channel permeation consistent with a gain-of-function phenotype.

**Figure 5:**
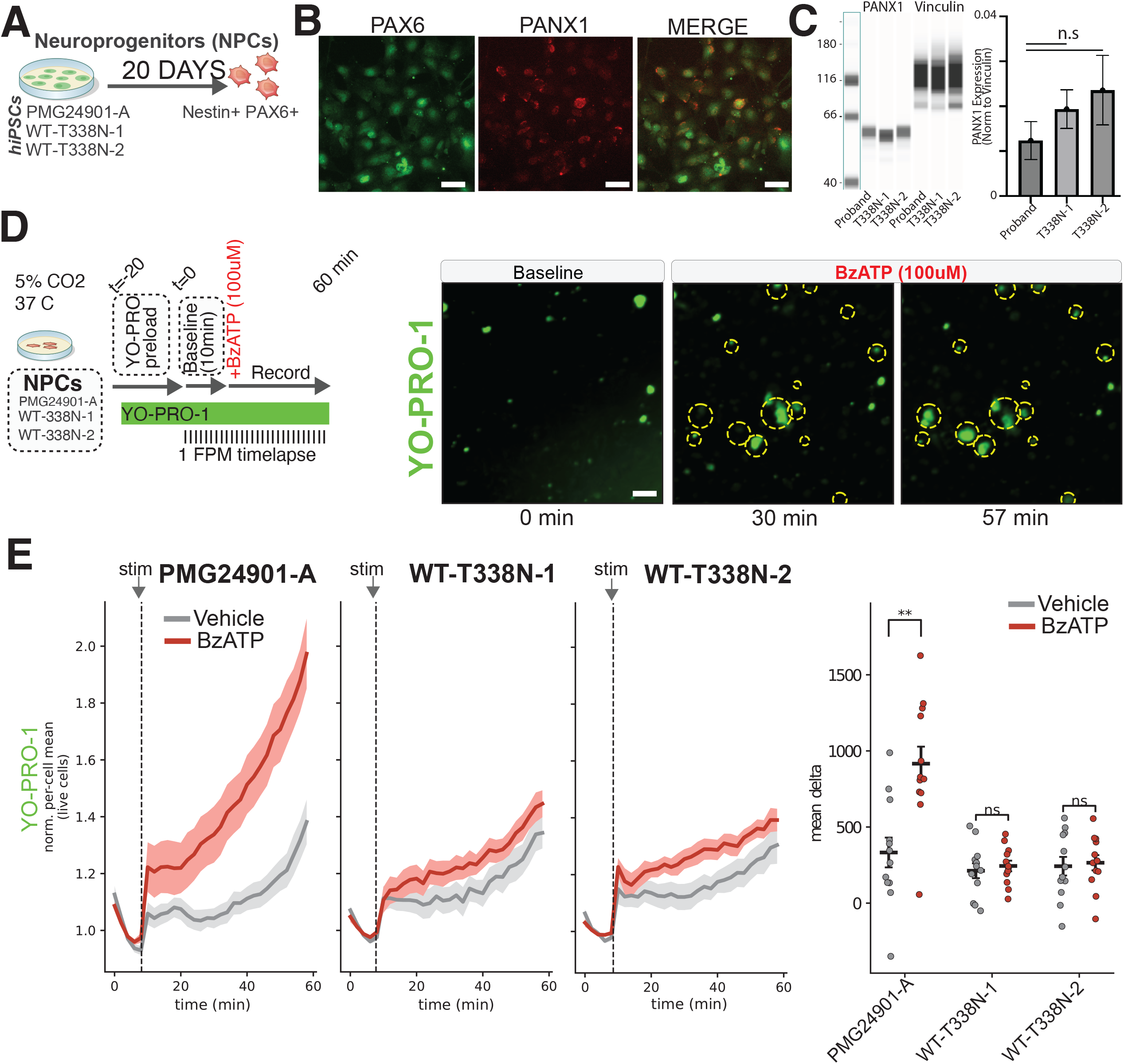
Functional characterization of PANX1 in proband-derived neural progenitor cells (NPCs). **(A)** Schematic illustrating 20-day differentiation protocol of hiPSCs into Nestin+/PAX6+ neural progenitors for the proband (PMG24901-A) and isogenic CRISPR corrected controls (WT). **(B)** Representative immunohistochemistry showing co-expression of PAX6 (green) and PANX1 (red) in the derived NPCs. Scale bar 20 μm. **(C)** Automated western blot (*left*) and quantification (*right*) of total PANX1 in proband and isogenic control NPCs normalized to Vinculin, demonstrating comparable expression across genotypes (one-way ANOVA; n.s. = not significant). Data from 4 independent NPC batches, presented as mean ± SEM. **(D)** *Left*, experimental timeline for the YO-PRO uptake timelapse assay. NPCs baseline fluorescence recorded for 10 minutes, followed by addition of 100 µM BzATP or vehicle, and recorded for an additional 60 minutes at 1 frame per minute (FPM). *Right,* representative fluorescent images of YO-PRO dye uptake (green) at baseline (0 min), 30 min, and 57 min post-BzATP. Dashed yellow circles highlight tracked cells actively taking up the dye. Scale bar 20 μm. **(E)** *(Left)* Traces of normalized per-cell mean YO-PRO fluorescence in live proband and isogenic NPCs. Shaded regions = standard error. *(Right)* Scatter plot quantifying the mean delta in fluorescence, demonstrating significant increase in dye uptake in PMG24901-A cells treated with BzATP compared to vehicle (p = 0.0011), which is absent in the isogenic controls (p = 0.72, WT-T338N-1; p = 0.86, WT-T338N-2; ns = not significant). Data points denote individual wells, 12 wells/line/condition. Significance determined by unpaired two-tailed Welch’s t-test.

## Discussion

Here, we demonstrate that activation of PANX1, a developmentally enriched ATP-releasing ion channel, can influence neurogenesis, cell fate, and cellular excitability in the developing cerebral cortex. We describe three unrelated individuals with de novo PANX1 missense mutations, PMG, and microcephaly, and demonstrate that these mutations disrupt complex glycosylation *in vitro* and confer gain-of-function effects on ATP release and ionic conductance. In mice and ferrets, *PANX1* overexpression causes cell-autonomous cell death and both cell-autonomous and non-cell-autonomous perturbations to neuronal migration, while proband-derived mutation-carrying neurons demonstrate increased stimulus-induced excitability, likely resulting from down-regulation of ion channel genes and neuronal differentiation pathways. Finally, mutant NPCs exhibit increased channel permeation, indicating that heightened ionic and metabolite flux is an early consequence of PANX1 disruption in the developing cortical column.

The clinical syndrome caused by the described *PANX1* missense mutations includes extensive and bilateral PMG, microcephaly, global developmental delay, and seizure disorders in two of three cases. In the gyrencephalic ferret, we found significantly reduced tangential spread of migrating electroporated cells in developing gyri and disrupted radial migration, mechanisms that could contribute to small gyri and cortical disorganization in humans. While we did not identify gross disruptions to gyral patterning at P10 following electroporation of WT or mutant *PANX1*, gyrification is still rudimentary at this age, and PANX1-induced cell death precluded analysis at P21, when gyral defects following IUE of other PMG-associated variants have been observed (28, 61, 62). The absence of detectable gyral defects may reflect cell death obscuring effects on affected gyri, electroporation of only a fraction of cells within a gyrus, or compensatory mechanisms between P10 and P21. In animal assays, we observed prominent cell death from mutant *PANX1* overexpression; affected individuals likely have some degree of cell death given that all probands are microcephalic, though less significant than expected from our overexpression findings. Additionally, because PANX1 forms a heptameric channel, how the proportion of WT and MT subunits in heterozygous individuals impacts channel function is unknown.

In this study, we demonstrate that PANX1 perturbation influences progenitor cell fate through a combination of *in vivo* and *in vitro* assays. In electroporated E17.5 mouse brains, both cell autonomous and non-cell autonomous disruptions to neuronal positioning were observed, suggesting fate changes. We further show that at P10 in mice, the proportion of layer 4 electroporated neurons is increased at the expense of layer II/III neurons, suggesting cell fate changes. The cellular positioning changes observed might be mediated by aberrant ionic flux in progenitors, as we observed in iPSC-derived NPCs. There may also be a role for gap junctions, resulting in non-cell-autonomous effects seen in electroporated mice. From transcriptomic analyses of NGN2 neurons, MT PANX1 was shown to drive vast gene regulatory changes indicative of altered neuronal cell state, with down-regulation of genes responsible for neuronal differentiation, and up-regulation of early developmental patterning genes. This may suggest a more immature cell state in the setting of aberrant PANX1 activity, which correlates with reduced dendritic branching observed (63). Taken together, our findings indicate that progenitors expressing MT *PANX1* have altered excitability and perturbed channel activity, generating widespread neuronal transcriptomic changes that alter cellular fate, which explains disrupted neuronal migration.

We found that both iPSC-derived NPCs harboring a PMG-associated *PANX1* mutation display gain-of-function (increased permeation) and increased excitability, consistent with seizures in two of three individuals in our cohort. Dysregulation of key ion channel genes and pathways suggests that excitability changes are not driven by PANX1 channel activity alone, but also by broader transcriptional changes, including altered expression of potassium channels such as KCNT1, which encodes the fetal-enriched KNa1.1 channel associated with a neurodevelopmental seizure disorder (64), and the recently identified epilepsy gene KCNJ4 (65). PANX1 can act as a signaling hub through channel-dependent and -independent mechanisms (34, 66, 67), potentially linking altered PANX1 function to these widespread transcriptional changes. Our RNA-seq data also support its established role in neuroinflammatory signaling through PANX1-mediated ATP release (68). Previous work has established that progenitor membrane potential regulates the timing of neurogenic divisions and temporal identity in the developing cortex (18), our findings suggest that aberrant *PANX1* may influence progenitor membrane properties through altered ion channel expression, thus regulating cell fate.

Previous investigations of gap junctions and hemichannels in early cortical development have centered on connexins (69–71), yet our work highlights a newfound role for pannexin hemichannels (and possible gap junctions (48, 72)) in cortical histogenesis. Our results implicate purinergic signaling and large pore channel physiology in the regulating membrane properties. Perturbation of PANX1 activity disrupted cortical progenitor identity, suggesting a role for paracrine signaling—or possibly electrical coupling via gap junctions—in progenitor cell cycle exit and migration. Furthermore, a pattern has emerged where malformation-associated ion channel variants are overwhelmingly gain-of-function, including activity-enhancing mutations in depolarizing channels (26–28), and activity-reducing or loss-of-function mutations in hyperpolarizing channels (73, 74). Together, these findings highlight how the balance and timing of ionic flux in development is essential for proper brain architecture.

## Methods

### Human subjects, clinical phenotyping, and genetics

The study was approved by the Boston Children’s Hospital institutional review board. Subjects were evaluated clinically, identified via the Matchmaker Exchange, and included with written informed consent. Clinical, radiographic, and genetic data were reviewed by neurologists, radiologists, genetic counselors, and at least two clinically trained study investigators. Detailed clinical descriptions are provided in Supplementary Table 1. De novo *PANX1* variants were identified and validated by research or clinical whole-exome sequencing as reported previously (25). Peripheral blood mononuclear cells from PMG24901 were used to generate iPSCs. De-identified postmortem human fetal brain tissue was obtained from the NIH NeuroBioBank at the University of Maryland under approved protocols and informed consent.

### Human developmental PANX1 expression analyses

BrainSpan/Allen Human Brain Atlas RNA-seq data (75) were used to examine *PANX1*, *PANX2*, and *PANX3* expression; neocortical RPKM values summarized to Gencode v10 exons were fit across age in R. *PANX1* expression was additionally assessed in 17-20 gestational-week human cortex by RNAscope (PANX1 and GFAP probes), immunostaining, and published MERFISH data (76). MERFISH analyses followed the published tissue-processing and imaging pipeline; cells below the tenth percentile of total transcript counts were filtered out and expression was normalized to total transcripts per cell. Additional details are provided in Supplementary Methods.

### PANX1 constructs and heterologous cell assays

Human PANX1-T2A-EGFP piggyBac constructs were generated in the iOn-CAG system using WT *PANX1* and site-directed mutagenesis to introduce *PANX1*-Asp14His, *PANX1*-Met37Arg, *PANX1*-Asn338Thr, or *PANX1*-Cys347His; plasmids were sequence verified. HEK293T and N2A cell transfections, ATP measurements, and western blotting are described in Supplementary Methods. Whole-cell voltage-clamp recordings were performed in transfected HEK293T cells using Sutter Instruments IPA. Current-voltage relationships were generated from -90 to +100 mV voltage ramps over 600 ms; capacitance and membrane resistance were calculated from negative current steps. Group comparisons used ANOVA with Tukey post hoc testing. Recording solutions and additional assay details are provided in Supplementary Methods.

### Mouse and ferret electroporation and tissue analysis

Animal experiments were approved by the Institutional Animal Care and Use Committee at Boston Children’s Hospital. CD1 mice of both sexes and ferrets (Mustela putorius furo) were used. Mouse in utero electroporation (IUE) was performed at E14.5 as described (77) using WT-*PANX1*, *PANX1*-Asn338Thr, or control transposon plasmids with piggyBase and pCAG-dsRed; embryos or pups were harvested at E17.5 or P10. For live migration imaging, brains were collected 48 h after IUE, sectioned at 300 µm, and imaged by spinning-disk confocal microscopy every 10 min for up to 24 h; cell trajectories were analyzed in Imaris.

Ferret electroporation followed established procedures (78, 79) and tissue was harvested at P1 or P21. Mouse and ferret brains were fixed in 4% PFA, cryoprotected, sectioned, and processed for fluorescence imaging or immunostaining. Confocal images were acquired on Zeiss systems and analyzed with Zen Blue and Fiji. Detailed electroporation parameters, tissue processing, and staining protocols are provided in Supplementary Methods.

### iPSC generation, correction, and NGN2 neuronal differentiation

PMG24901 iPSCs generation and characterization is described in Supplementary Methods. Two isogenic WT lines were generated by CRISPR-Cas9-mediated correction of *PANX1* Thr338Asn and clonal genotyping. Feeder-free iPSCs were maintained in mTeSR Plus and routinely monitored for mycoplasma and karyotype abnormalities.

Forebrain glutamatergic neurons were generated by lentiviral TetO-Ngn2-Puro/FUW-M2rtTA transduction followed by doxycycline induction, puromycin selection, neurotrophic support, and Ara-C treatment. On differentiation day 6, neurons were dissociated and replated with human astrocytes for electrophysiology/MEA studies or onto coated 96-well plates for morphometric analysis. Three independent differentiation batches were performed with all cell lines. Full induction, plating, and maintenance conditions are provided in Supplementary Methods.

### Neuronal morphology, electrophysiology and multielectrode array

For neurite analysis, day-6 NGN2 neurons were replated at 40,000 cells/well, fixed after 24 h, immunostained for MAP2 and Hoechst, and imaged on an ImageXpress MicroXLS microscope. Eight wells per line and nine fields per well were analyzed in each batch using the MetaXpress neurite algorithm. Whole-cell patch-clamp recordings were performed on DPD 26-32 using SutterPatch IPA. Current- and voltage-clamp protocols assessed action-potential properties, excitability, membrane resistance, capacitance, and ionic currents; recording solutions provided in Supplementary Methods. For MEA, NGN2 neurons were plated with human astrocytes on 48-well CytoView MEA plates and spontaneous network activity was recorded for 25 d using the Maestro system (Axion Biosystems). Ten-minute recordings were acquired at 12.5 kHz after 10 min acclimation and analyzed in AxIS Navigator. Weighted mean firing rate and multi-electrode simultaneity were quantified; parameter trajectories were fit to three-parameter sigmoidal curves and coefficients compared across genotypes in R.

### Bulk RNA extraction, sequencing and bioinformatic analysis

RNA was isolated from monolayer cultures by TRIzol/1-bromo-3-chloropropane phase separation and isopropanol precipitation and sent for bulk RNA sequencing. Differential expression was analyzed in R (v4.4.0) using DESeq2 (v1.46.0) (60), with genotype as the main effect and sequencing run and read depth included as batch covariates. Genes with <5 reads per sample were removed; significance thresholds were adjusted p < 0.01 and |log2 fold change| >2. EnhancedVolcano (v1.24.0) (80) was used for visualization. Up- and down-regulated genes were analyzed separately in Metascape (v3.5.20260701) (81); terms with log q-value < -1.0 are reported in Supplementary Dataset 2.

### iPSC-derived NPCs and YO-PRO-1 assay

NPC generation and immunocytochemistry is described in Supplementary Methods. PANX1 channel activity was measured by YO-PRO-1 uptake. Patient and isogenic NPCs were loaded with 10 µM YO-PRO-1 in Ca2+/Mg2+-free HBSS with 20 mM HEPES and imaged on an Incucyte system at 1 frame/min. After a 10-min baseline, cells received 500 µM BzATP or vehicle and were imaged for 50 min. Per-cell fluorescence was quantified from segmented, drift-corrected images using a custom Python pipeline; well-level change in fluorescence was calculated relative to baseline and BzATP versus vehicle responses compared by Welch’s t test.

### Quantification and statistical analysis

Statistical analyses were performed in Prism unless otherwise specified, with individual tests reported in figure legends. Significance was defined as p < 0.05 unless otherwise specified, and bar-plot error bars represent SEM. Data normality was assessed by examining distribution and spread of data.

## Acknowledgments

We are grateful to the participant families involved in this study, and we thank the referring providers of PMG24901 and PMGSL101, Andrea V. Murphy, Kara C. Klemp, and Stephen R. Braddock. We are grateful to Drs. Annapurna Poduri, Lisa Goodrich, Chinfei Chen, Leigh Anne Swayne, Leigh Wicki-Stordeur and Gary Yellen for helpful discussions. This research was conducted with support from the Human Neuron Core within the Rosamund Stone Zander Translational Neuroscience Center, Boston Children’s Hospital, supported by IDDRC (NIH P50HD105351). iPSC lines were generated and characterized by the Harvard Stem Cell Institute core facility. We thank J.E.N. and the NIH Neurobiobank at the University of Maryland Brain and Tissue Bank for facilitating tissue collection. N.K.H. is supported by NINDS 1F31NS134253-01A1, the Albert J. Ryan Foundation, and is a Stuart H.Q. and Victoria Quan Fellow in the Harvard Department of Neurobiology. N.K.H. and M.T are supported by NIH T32 GM007753 and T32 AG000222; D.E-A. by EMBO Postdoctoral Fellowship ALTF 336-2022; R.E.A. by the Autism Speaks Postdoctoral Fellowship 13008 and NIGMS T32 GM007748; T.M.B. by the Harvard/MIT Equitable Access to Research Training (HEART) MD–PhD Summer Program at Harvard Medical School; X.Q. by the NINDS (R00 NS135123). C.A.W. is supported by NINDS (R01 NS035129) and R.S.S. by NIH R00 NS112604, R01NS140046, and DP2NS148744. C.A.W. is supported by the Allen Discovery Center for Human Brain Evolution through The Paul G. Allen Frontiers Group. C.A.W. is an Investigator of the Howard Hughes Medical Institute.

## Author Contributions

N.K.H., R.S.S., and C.A.W. designed research; N.K.H., D.J.K., S.C.D., D.E.-A., C.N.C., S.R.G., M.B., Q.C., K.I.S., J.E.N., E.M.D., T.M.B., S.R.B., and S.K.A. investigated; N.K.H, D.E.-A., R.E.A., M.T., C.H., R.H. and S.M. contributed new reagents/analytic tools; N.K.H., D.J.K., S.C.D., C.N.C., S.R.G., X.G.M., K.I.S., J.E.N., T.M.B, X.Q., A.D.D., M.T., C.C., and L.S. analyzed data; S.S., C.A.W. and R.S.S. supervised research; C.A.W. and R.S.S. edited manuscript; and N.K.H. wrote the paper.

