## Supplementary Information for "Pannexin-1 regulates cellular identity and excitability in the developing cortex"

**This PDF file includes:**

Supplementary Discussion
Figures S1 to S15
Tables S1 to S2, and legends for Supplementary Datasets 1 and 2
Legends for Movies S1 to S3
Supplementary Methods
SI References

**Other supporting materials for this manuscript include the following:**

Movies S1 to S3
Supplementary Datasets 1 and 2

**Supplementary Discussion**

PANX1 effects on ionic and metabolite flux observed in this study may be explained by electrostatic changes introduced by the mutations, leading to disrupted channel gating or ion permeation. D14 is located within the N terminal domain (NTD) and may alter its movement into and out of the pore. In structures of PANX1 with the NTD flipped down (associated with no activity, or a closed state), there is an accumulation of negative charges near the narrowest point of the permeation pathway (1). This charge accumulation was proposed to serve as an electrostatic barrier to anion permeation (1). D14H replaces a negative charge with a more positively charged amino acid, which may hinder movement of the NTD away from this electrostatic barrier. M37R, nearby to the NTD, replaces a hydrophobic side chain with a positively charged one, further suggesting altered electrostatic potential as a possible mechanism of enhanced channel activity. N338 is located just upstream of the C terminal activating domain (CAD, between residues I361 and L370) (1). This domain promotes channel opening and facilitates NTD movement (1). Considering N338T, both asparagine and threonine are polar, but the hydroxyl group on threonine contributes to greater polarity, which may strengthen CAD interactions and contribute to channel opening. Analyses such as single particle cryo-EM to resolve channels containing each mutant are needed to definitively assess how they might alter structural properties.

With respect to our RNA sequencing data in NGN2 neurons, our results confirm a role in neuronal development, cell fate as well potential cell stress activation, it’s noteworthy the ectopic expression of GABAergic-associated genes like GAD1, which was upregulated by the proband variant. NGN2-induced neurons can give rise to heterogeneous central and peripheral neuronal populations; however, these subpopulations are excitatory in nature with no evidence of generating GABAergic subpopulations (2). The lack of GABAergic enrichment suggests that the affected genotype might not necessarily be driving a GABAergic fate, but rather ectopic GAD1 expression could be a secondary effect of the upregulation of multiple transcription factors associated with pattern specification and cell fate, some of which are also upstream of inhibitory (FOXN4, DLX1 DLX2, DLX5 and DLX6) lineages (3). While there were also dysregulated transcription factors from excitatory fate (TBR1, FEZF2), specification and cell identity programs in other structures like the thyroid (PAX8) (4) were also ectopically expressed.

Gain of function variants in *PANX1* have been previously identified in women with infertility (5-9). Many mutations were paternally inherited, and those that were characterized disrupted complex glycosylation, and increased channel conductance and ATP release, a match to the cellular phenotype we describe here. Notably, none of the reports connecting *PANX1* to infertility discuss co-morbid findings of cortical malformations, microcephaly, or seizures. Given the pediatric presentation of such features in our cases with *de novo* mutations, the PMG-associated alleles are likely more developmentally disruptive as it is improbable that the cohort of women with *PANX1*-associated infertility have severe, undiagnosed neurologic disease. However, it is possible to have subclinical cortical malformations with minimal impact on cognition that could go undiagnosed without an indication for MRI (10). This raises the possibility of a spectrum of disease caused by *PANX1* variants, with infertility alone as a milder presentation of channel hyperactivity and cell death. It is unknown whether the pediatric cases we present (two females and one male) might ultimately have infertility. Further investigations to elucidate the mechanisms of PANX1-related diseases and to discern the genotype-phenotype correlations can aid in the diagnosis and management of patients.

The results of *PANX1* loss of function in cortical development are not completely clear. A single individual with a multisystem disorder was identified to harbor a homozygous *PANX1* variant that reduces channel function (11); however, additional genetic matches to this syndrome have not yet been reported. In addition, the individual harbors additional homozygous and heterozygous gene variants that are possibly pathogenic and were not functionally validated (11). Knockout (KO) of PANX1 in human organoids resulted in significantly smaller organoids, with gene expression changes related to WNT signaling and cell adhesion (12), consistent with essential roles in neurogenesis. However, *Panx1* KO mouse brains are grossly normal (13, 14), although their layer 5 neurons have reduced dendritic spine densities (15). In previous studies, blocking PANX1 or KO in mice has been shown to reduce neuronal discharges and epileptiform activity (16-19), and can spare cortical tissue in female mice following cerebral artery infarct (20).

Together these results suggest that while loss of *PANX1* in mice is non-essential for cortical structure, it causes other changes in physiology, and human-specific essential roles for *PANX1* may yet come to light.

**Limitations**

In this study, we performed *in utero* electroporations in mice and ferrets to assess developmental phenotypes related to *PANX1* perturbation. However, because this approach relies on exogenous plasmid expression, we cannot exclude that overexpression may alter cellular physiology and induce non-native toxicity. Consequently, conclusions related to neuronal positioning and cell death may partially reflect supraphysiologic levels of *PANX1* expression. To mitigate this limitation, our study heavily incorporates iPSC-derived neurons and NPCs, which provide a robust complementary model system featuring endogenous expression in the context of the patient genotype. Future *in vivo* studies utilizing gene-editing strategies in mice or ferrets will be valuable to assess *PANX1* perturbation under physiologic expression levels.

### Supplementary Figures

**A**

**PMG20601:** c.40G>C, Asp14His

| | |
|---|---|
| **Human** | AQLATEYVFSDFLLKEPTEPK |
| **Chimpanzee** | AHLATEYVFSDFLLKEPTEPK |
| **R. Macaque** | AHLATEYVFSDFLLKEPTEPK |
| **Mouse** | AHLATEYVFSDFLLKEPTEPK |
| **Rat** | AHLATEYVFSDFLLKEPTEPK |
| **Cat** | AHLATEYVFSDFLLKEPSEPK |
| **Ferret** | AHLATEYVFSDFLLKEPSEPK |
| **Opossum** | AHLATEYVFSDFLLKEPSEPK |
| **Chicken** | AHIATEYVFSDFLLKEPPETR |
| **Sea Turtle** | ARTAAEYMLSDALLPDPNGSR |
| **Xenopus** | AHIATEYVFSDFLLKDPPESK |
| **Zebrafish** | AHIATEYVFSDFLLKDPPESK |
| **Lamprey** | AAAAAEYIFSDSLLKDSADAR |

**PMGSL101:** c.110T>G, Met37Arg

| | |
|---|---|
| **Human** | GLRLELAVDKMVTCIAVGLPL |
| **Chimpanzee** | GLRLELAVDKMVTCIAVGLPL |
| **R. Macaque** | GLRLELAVDKMVTCIAVGLPL |
| **Mouse** | GLRLELAVDKMVTCIAVGLPL |
| **Rat** | GLRLELAVDKMVTCIAVGLPL |
| **Cat** | GLRLELAVDKMVTCIAVGLPL |
| **Ferret** | GLRLELAVDKMVTCIAVGLPL |
| **Opossum** | GLRLELAVDKMVTCIAVGLPL |
| **Chicken** | GLRLELALDKIVTCIAVGLPL |
| **Sea Turtle** | GLRLELPSDRVVKFVTVGLPL |
| **Xenopus** | GLRLELAVDKLVSCIAVGLPL |
| **Zebrafish** | GIRLDLALDKIVTCVAVGLPL |
| **Lamprey** | GLRLELPADRLLKFITVGLPL |

**PMG24901:** c.1013A>C, Asn338Thr

| | |
|---|---|
| **Human** | LSLYNLFLEENISEVKSYKCL |
| **Chimpanzee** | LSLYNLFLEENISEVKSYKCL |
| **R. Macaque** | LSLYNLFLEENISEVKSYKCL |
| **Mouse** | LSLYNLFLEENISELKSYKCL |
| **Rat** | LSLYNLFLEENISELKSYKCL |
| **Cat** | LSLYNLFLEENISELKSYKCL |
| **Ferret** | LSLYNLFLEENVSELKSYRCL |
| **Opossum** | LSLYYLFLEENISELKSYKCL |
| **Chicken** | LSIYLLFLEENVSELKSYKCL |
| **Sea Turtle** | LSLYLLFIEENVSELKSYKCL |
| **Xenopus** | HSLFLLFLEENVSELKSYKFL |
| **Zebrafish** | LALYLLFLDENLSELKSYKCL |
| **Lamprey** | - - - - - - - - - - - - - - - - - - - - - |

**B**

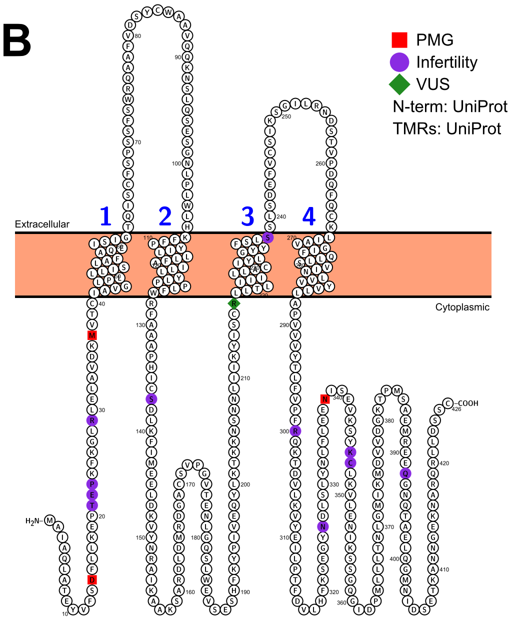


**Figure S1: Evolutionary conservation of *PANX1* PMG variants. (A)** Each proband variant (marked in red) occurs at an amino acid highly conserved among mammals. Asp14 and Asn338 are amino acid residues conserved in vertebrates. Lamprey lacks the C terminal region of *PANX1*. Residue changes at nearby amino acids marked in blue. **(B)** PMG associated mutations in *PANX1* highlighted in linear protein structure in red compared to previously reported mutations in *PANX1*. Rendered in protter.com. See Table S2.

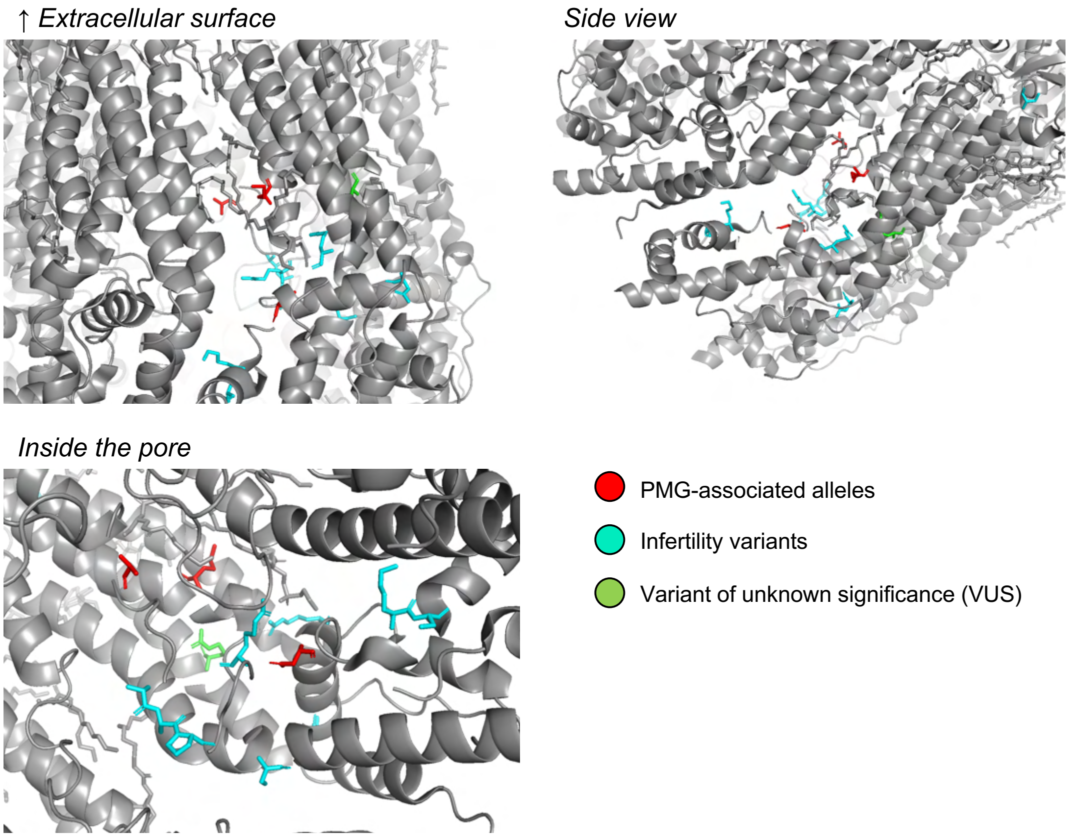


**Figure S2: Previously reported *PANX1* disease-associated residues in relation to PMG-associated residues.** Heptameric cryo-EM structure 6WBF of PANX1 from Ref. (21) rendered in PyMol. In a single subunit, three PMG-associated amino acid residues are highlighted in red (Asp14, Met27, and Asn338), along with a single variant of unknown significance (VUS, green) and 8 out of 9 total variants identified in women with infertility (Thr21-Pro23, Arg29, Ser137, Ser238, Arg300, Asn326, Lys346, and Cys347; cyan) (5-9). See Table S2.

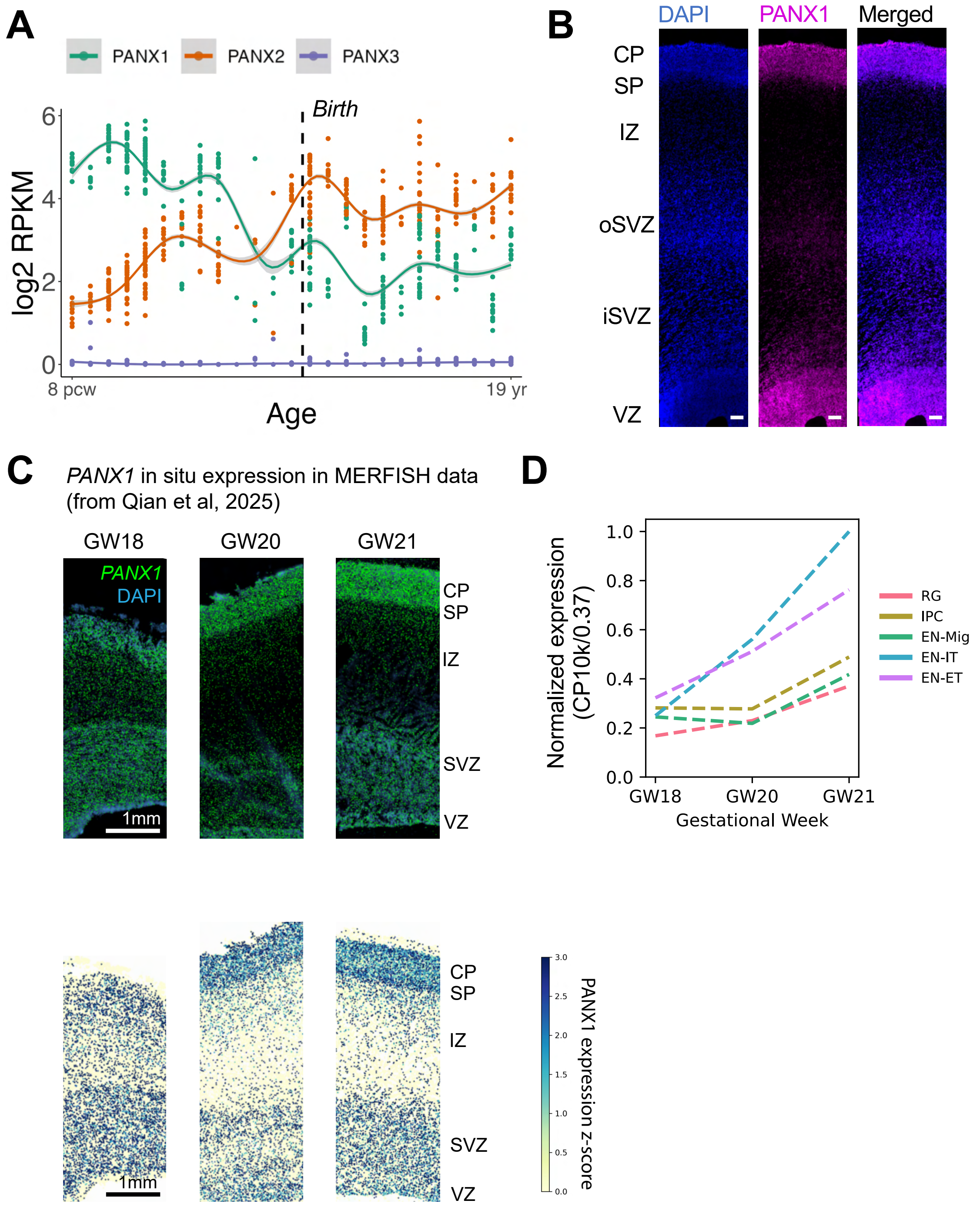


**Figure S3: RNA expression of PANX1 in the developing human cortex. (A)** Bulk RNA sequencing of human cortex from 8 post-conception weeks (pcw) to 19 years old demonstrates temporal enrichment of *PANX1* expression in early gestational development, with decreasing expression just before birth, around the end of gyrification. *PANX2* expression is reduced gestationally relative to its postnatal expression, demonstrating the opposite expression profile to *PANX1*. *PANX3* transcripts were lowly detected in cortical tissue, consistent with its lack of expression in the brain. Analysis of data obtained from the publicly available Allen Developing Human Brain Atlas. **(B)** RNAscope in a 19 gestational week fetal cortex demonstrates *PANX1* expression in both the VZ/SVZ and cortical plate. Scale bar = 200μm. VZ, ventricular zone; iSVZ, inner subventricular zone; oSVZ, outer SVZ; IZ, intermediate zone; SP, subplate; CP, cortical plate. **(C)** MERFISH *PANX1* in situ expression from three gestational ages; analysis of data obtained from Qian et al (22). **(D)** Quantification of normalized *PANX1* expression across cell types over GW18, 20, and 21, suggesting increasing RNA expression between GW18 and 21; analysis of data obtained from Qian et al (22).

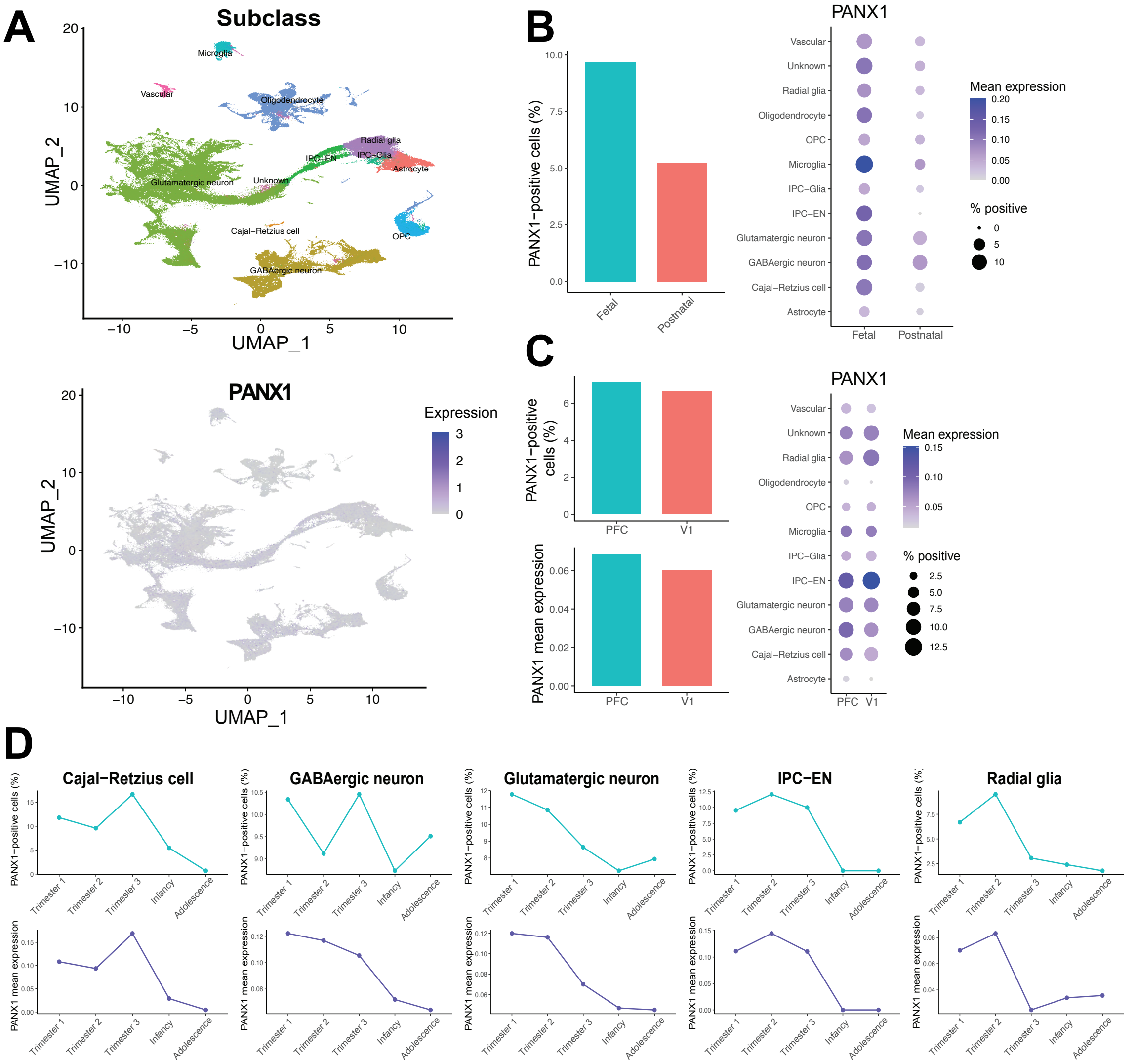


**Figure S4: Single cell RNA-seq expression of *PANX1* in human brain development. (A)** UMAP and feature plots of single cell RNA expression of *PANX1* in human developmental cell types analyzed from Wang et al (23). **(B)** *Left:* Percent of *PANX1*+ cells is enriched during fetal timepoints compared to postnatal; fetal = first, second, and third trimesters, postnatal = infancy and adolescence. *Right: PANX1* is broadly expressed across multiple cell types, with modest enrichment in neurons and microglia. OPC = oligodendrocyte precursor cell; IPC = intermediate progenitor cell. **(C)** *Left: PANX1* expression is comparable between prefrontal cortex (PFC) and area V1. *Right:* cell-type specific expression of *PANX1* between PFC and V1 shows similar levels of cellular expression across regions. **(D)** Levels of *PANX1* expression in Cajal-Retzius cells, intermediate progenitor subtypes, radial glial cells, and neurons demonstrate a declining trend between first trimester and infancy.

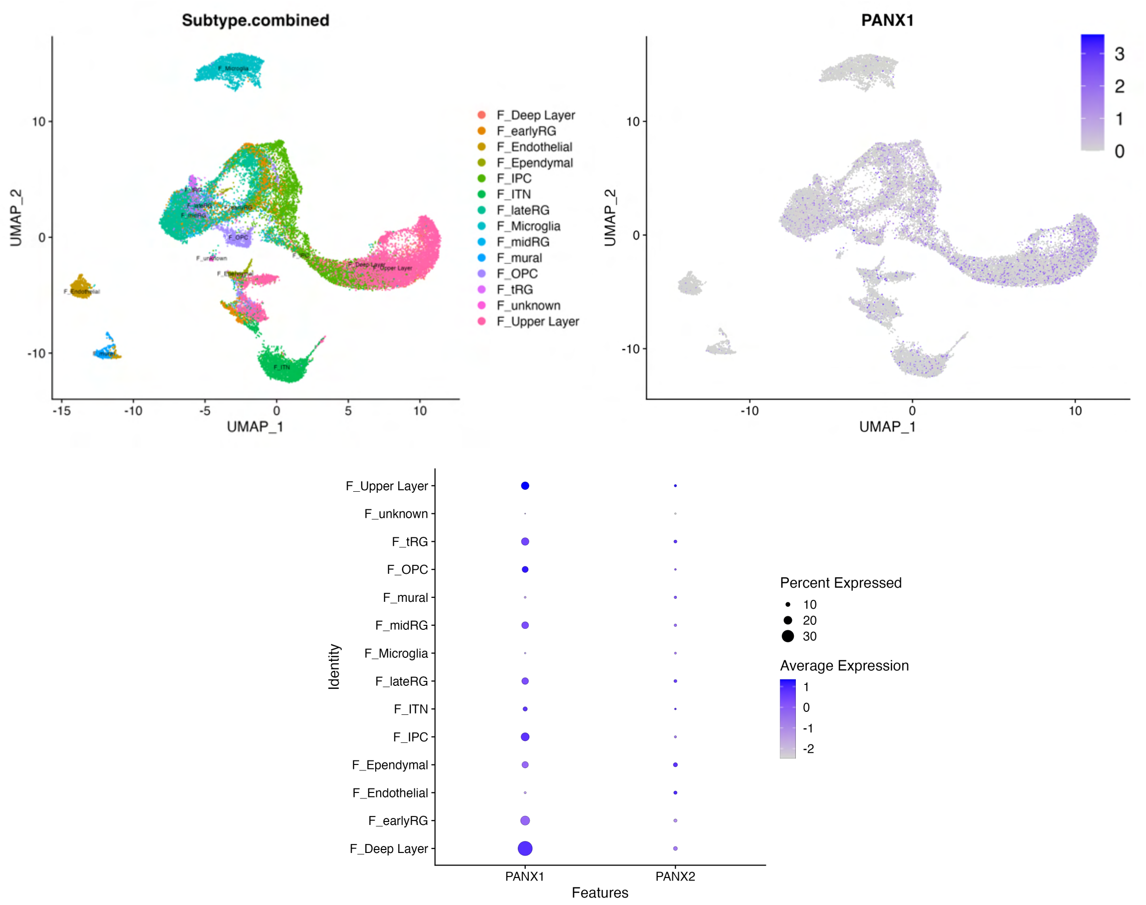


**Figure S5: Single cell RNA expression of *PANX1* in ferret cortex.** *Top*, UMAP plot of single cell expression of PANX1 in ferret cortex during corticogenesis from six developmental timepoints (embryonic day E25, E34, E40, and postnatal day P5 and P10) (Bilgic et al. (24)). PANX1 is diffusely expressed across multiple cell types in development. *Bottom*, dot plot reveals enrichment of *PANX1* in neurons, glia and progenitors, with some enrichment in deep layer neurons. Within RG cells, there is a mild enrichment in early RG and tRG cells compared to late RG cells, suggestive of an early role in progenitor specification. RG = radial glia, tRG = truncated RG, ITN = interneurons, IPC, intermediate progenitor cell, OPC = oligodendrocyte precursor cell.

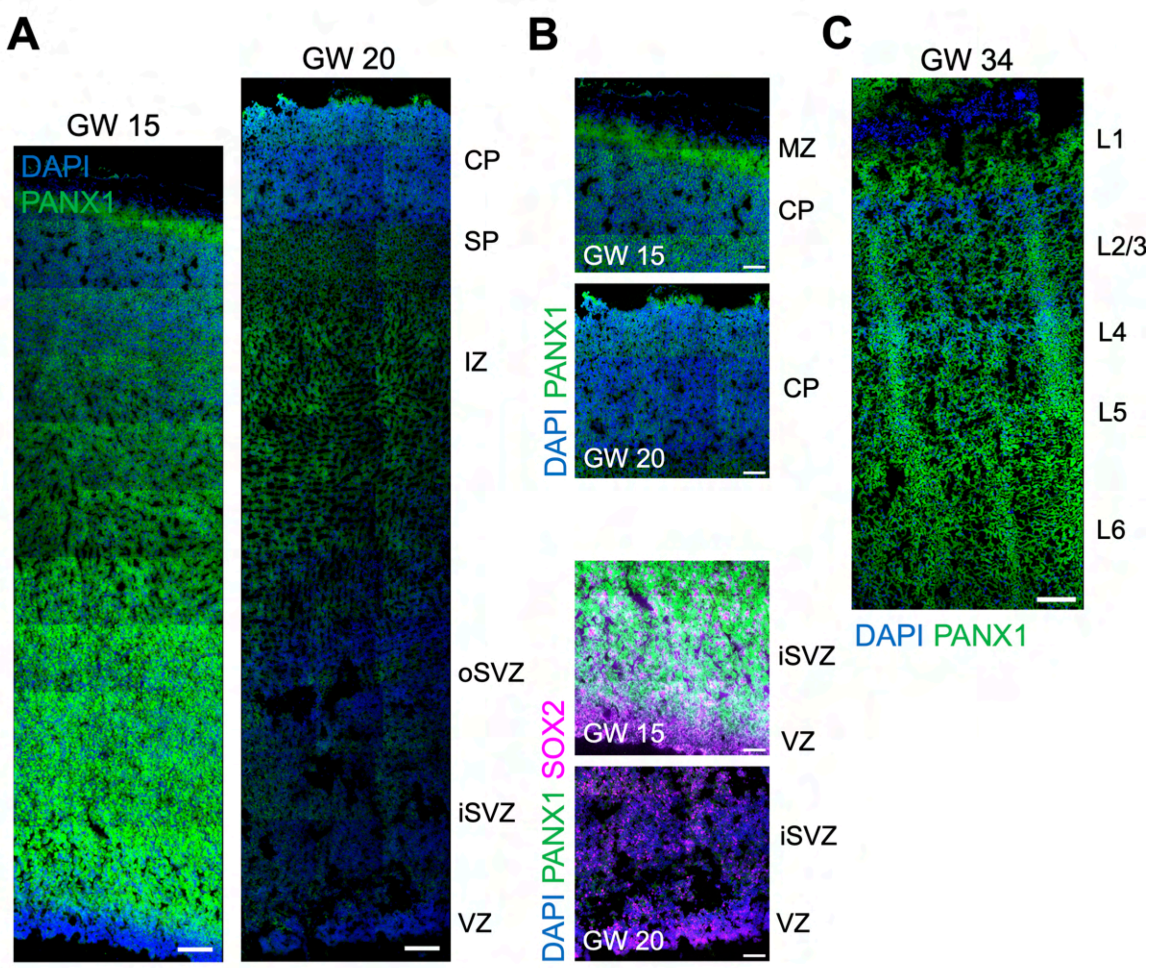


**Figure S6: PANX1 immunoreactivity is enriched in early gestational development. (A)** PANX1 immunoreactivity in two fetal cortex samples, at 15 and 20 gestational weeks with increased expression at GW15. Scale bar = 200μm. **(B)** Zoom in of the CP and VZ/SVZ for both fetal samples. Scale bar = 100μm. **(C)** PANX1 expression at GW 34 across the CP. Scale bar = 200μm.

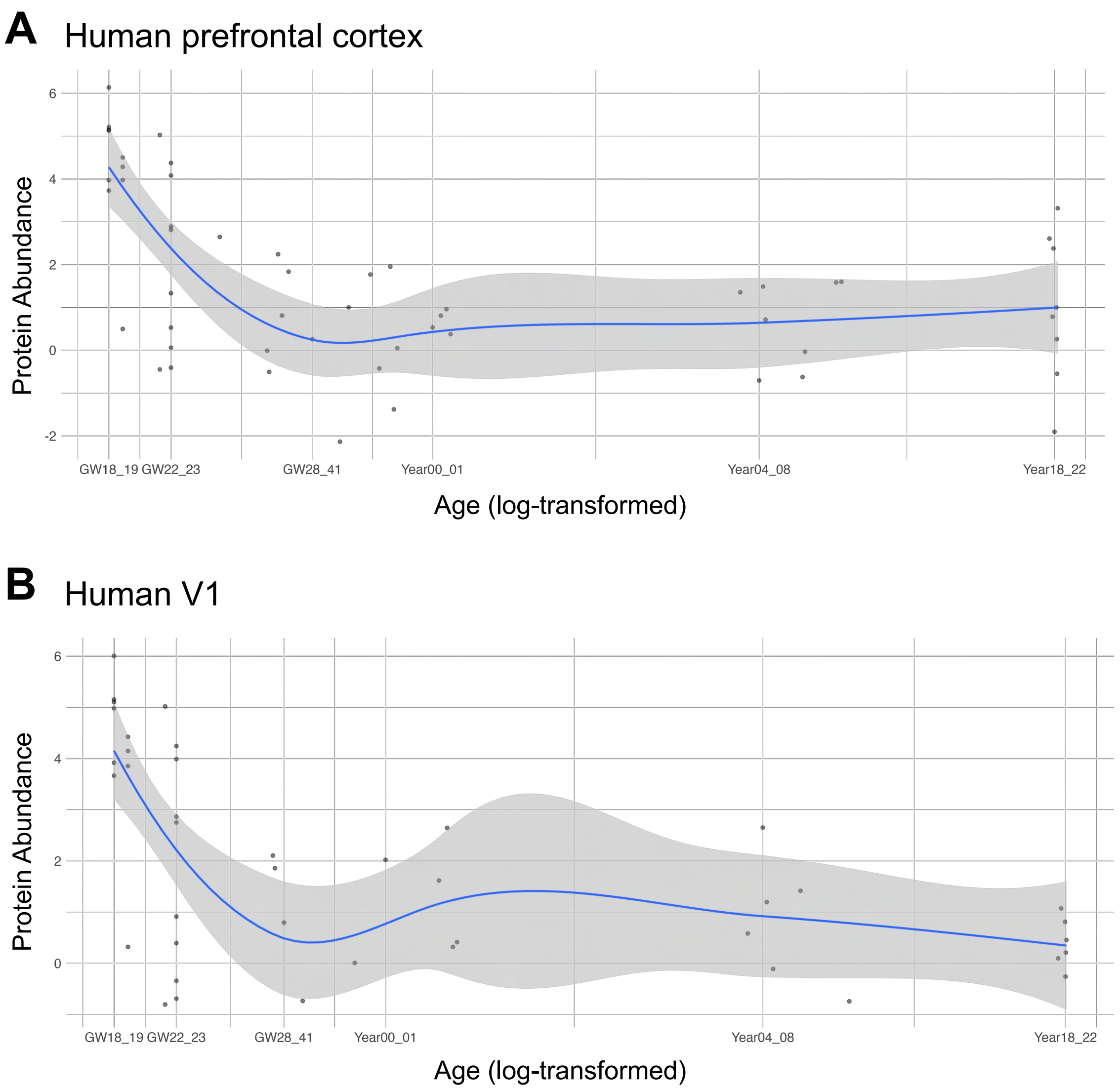


**Figure S7: Expression of PANX1 in human synaptosomes throughout development. (A)** Protein expression of PANX1 in human prefrontal cortex obtained from human post synaptic densities (PSDs) reveals enriched PANX1 expression in early gestational development. **(B)** PANX1 PSD expression in human visual cortex recapitulates gestational enrichment of PANX1. Data in **(A)** and **(B)** obtained from the interactive portal https://liwang.shinyapps.io/PSD_development_explorer/ and reported in Ref (25).

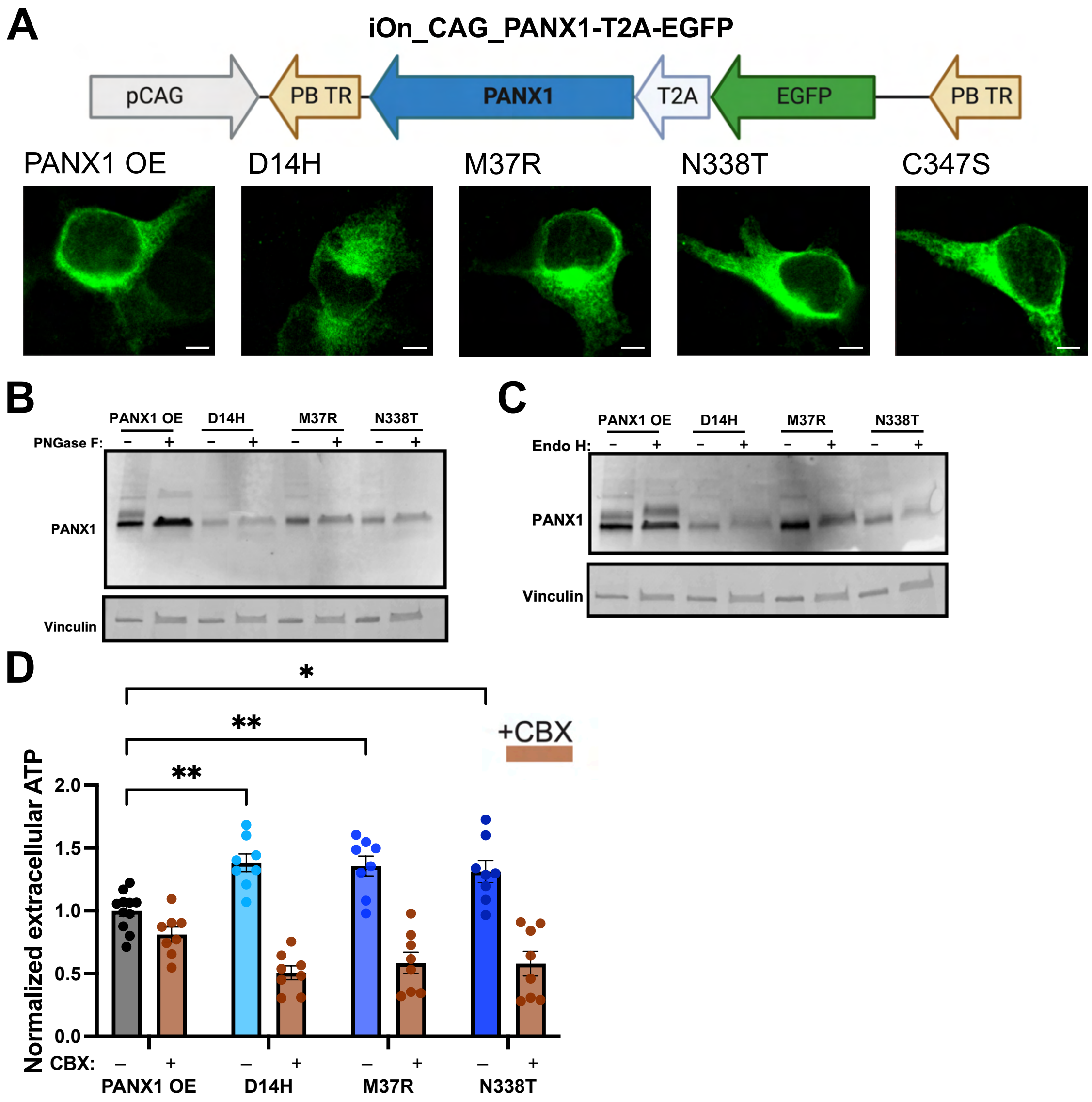


**Figure S8: *PANX1* mutations disrupt glycosylation and increase ATP release—related to Figure 1. (A)** *Top*, schematic of iOn_CAG-PANX1-EGFP. Driven by a pCAG promoter, a transposon contains either WT or mutated PANX1 (D14H, M37R, N338T, or C347S) and EGFP linked by a T2A peptide to obtain their separate expression. The transposon is inserted between piggyBac recognition sites in parallel orientation, limiting the expression of this construct based on its integration into a host cell (26). *Bottom*, representative images of HEK293T cells transfected with integrating piggyBac *PANX1* constructs immunostained for PANX1. Transfections demonstrate no difference in the localization of PANX1 expression. Scale bar = 5μm. PB TR, piggyBac terminal repeat. **(B, C)** Additional western blotting for PANX1 demonstrates specificity of bands for glycosylated species Gly0, Gly1, and Gly2. **(B)** PNGase F enzymatic digestion of HEK293T cell protein lysate transfected with either WT or mutant PANX1. Loading control of Vinculin in the bottom panel. **(C)** Endo H enzymatic digestion of protein lysate of transfected HEK293T cells. **(D)** In transfected N2A cells, PANX1 mutants demonstrate increased ATP release relative to WT-transfected cells, as quantified using the ATP Determination Kit (two-way ANOVA with Sidak's multiple comparisons test, * $p<0.05$, ** $p<0.01$). ATP values for each transfection are normalized to the amount of protein per well. All values are additionally normalized to the average extracellular ATP from WT conditions. Addition of CBX (brown bars) inhibits ATP release and normalizes values from mutant conditions to that of the WT condition (no significant difference, two-way ANOVA with Sidak's multiple comparisons test).

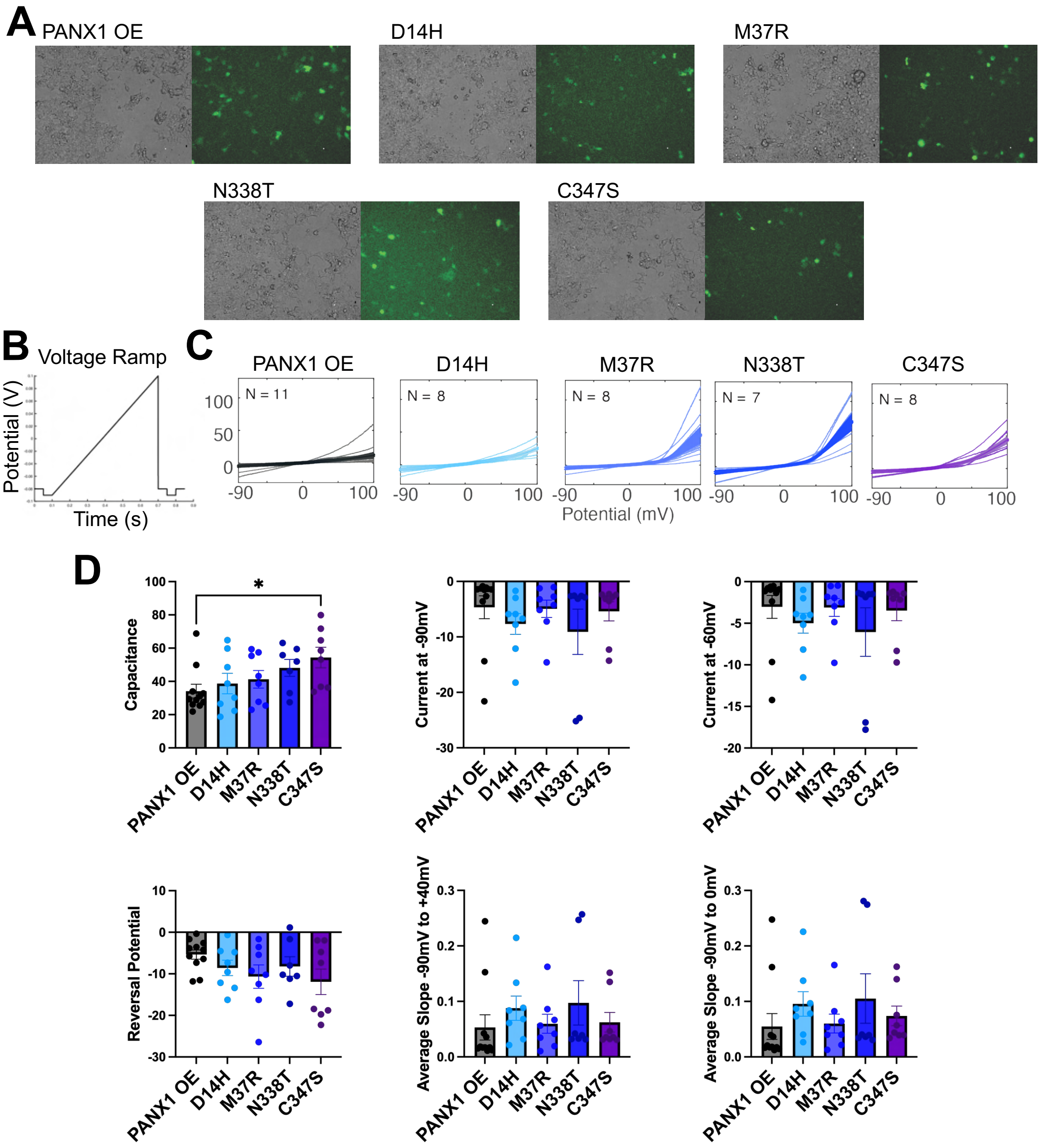


**Figure S9: Whole cell voltage-clamp recording in transfected HEK293T cells. (A)** Example images of 293T cells transfected with either WT or mutant, MT, protein with brightfield (*left*) and EGFP expression (*right*). **(B)** Voltage-clamp ramp protocol. **(C)** Individual ramps recorded for replicates of each condition. **(D)** Additional parameters recorded from whole-cell voltage clamp recordings. The overall capacitance of cells transfected with Cys347Ser were slightly increased compared to WT ($p = 0.271$, one-way ANOVA with multiple comparisons test); measurements of current density were normalized to the capacitance for each replicate. No significant differences in the current at -90mV, -60mV, the reversal potential, or the average slope from -90mV to 0mV or +40mV ($p > 0.05$, one-way ANOVAs with multiple comparisons test).

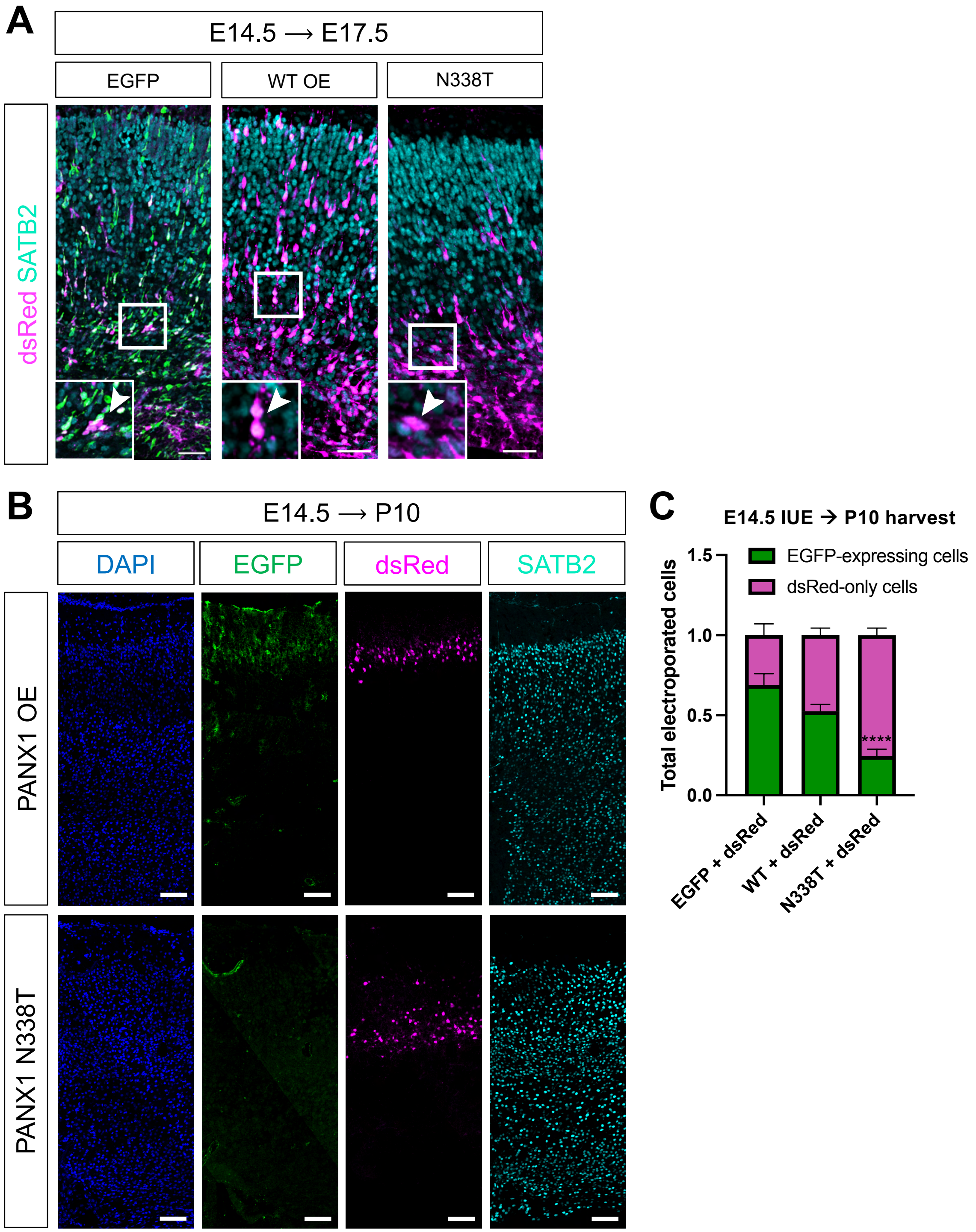


**Figure S10: Overexpression of *PANX1* disrupts positioning of neuronal cells across development in mice. (A)** SATB2+ IUE cells at E17.5 localize predominantly to the VZ and SVZ of N338T-electroporated mice. Scale bar = 50µm. **(B)** Representative immunofluorescence staining of mouse P10 cortex demonstrates the distribution of SATB2+ IUE cells in both WT and N338T conditions (scale bar = 100µm). **(C)** At P10, there is a significant reduction in EGFP-expressing electroporated cells in N338T-electroporated mouse brains compared to EGFP control + dsRed ($p < 0.0001$), using two-way ANOVA with Sidak multiple comparisons test.

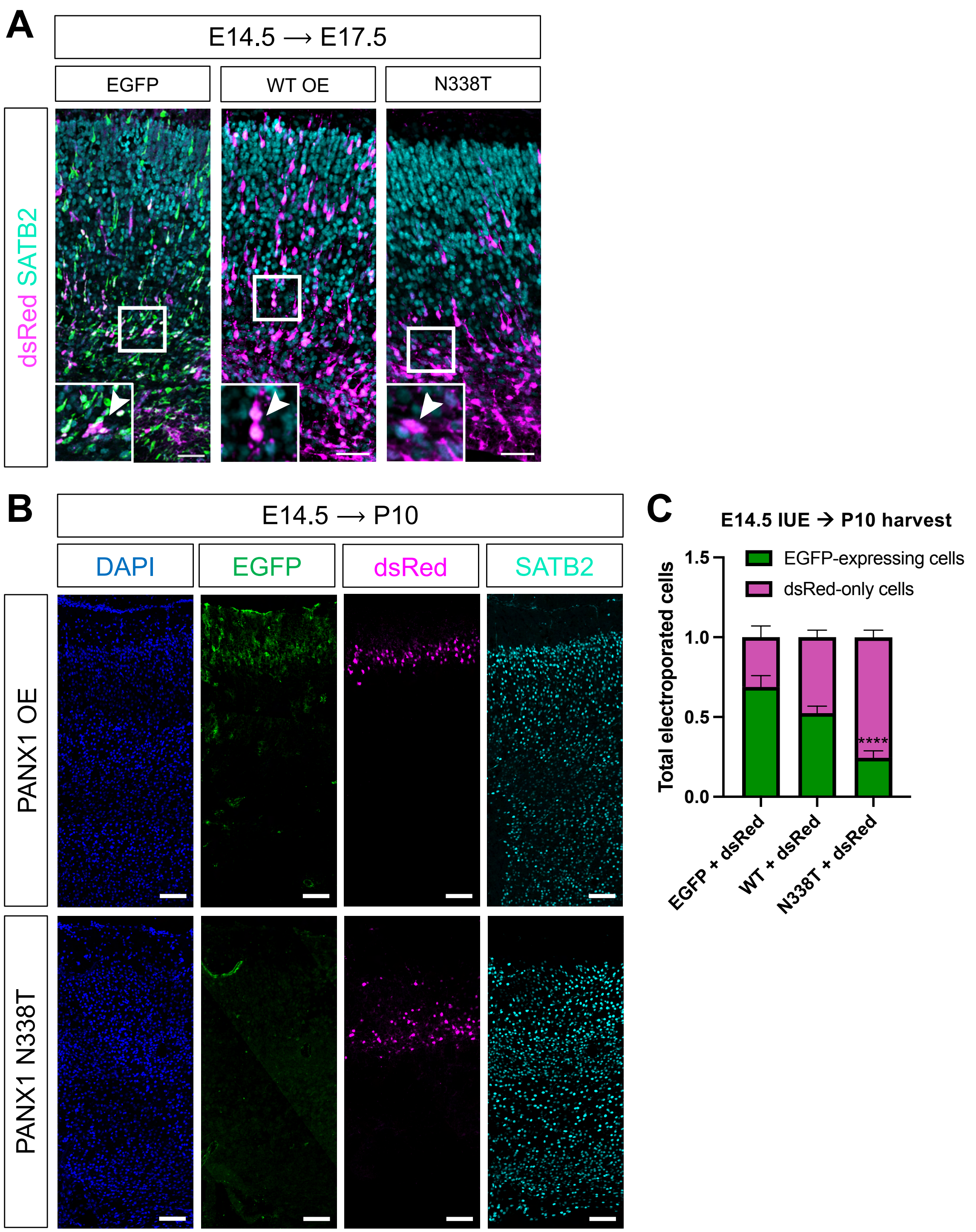


**Figure S11: Post-natal electroporation of ferret visual cortex and CC3 analysis. (A)** A total of 5 P1 kits were electroporated with either WT or MT PANX1 and harvested at P10 (scale bar = 3000μm). Mutant P10 brains reveal no gyral abnormalities in the visual cortex. All electroporations targeted the right hemisphere for both conditions. **(B)** Representative confocal images of electroporated P10 ferret visual cortex stained for the apoptotic marker CC3. **(C)** 25-35% of EGFP+ electroporated cells stain positively for CC3, with a slight increase in the proportion of apoptotic electroporated cells in N338T mice compared to WT (p = 0.1086, t-test, n

= 5 per condition, scale bars = 200μm). **(D)** Example of ferret visual cortex electroporation visualizing both EGFP+ and dsRed+ electroporated cells. Staining for neuronal markers demonstrates migrating cell types are SATB2+ (scale bar = 200μm).

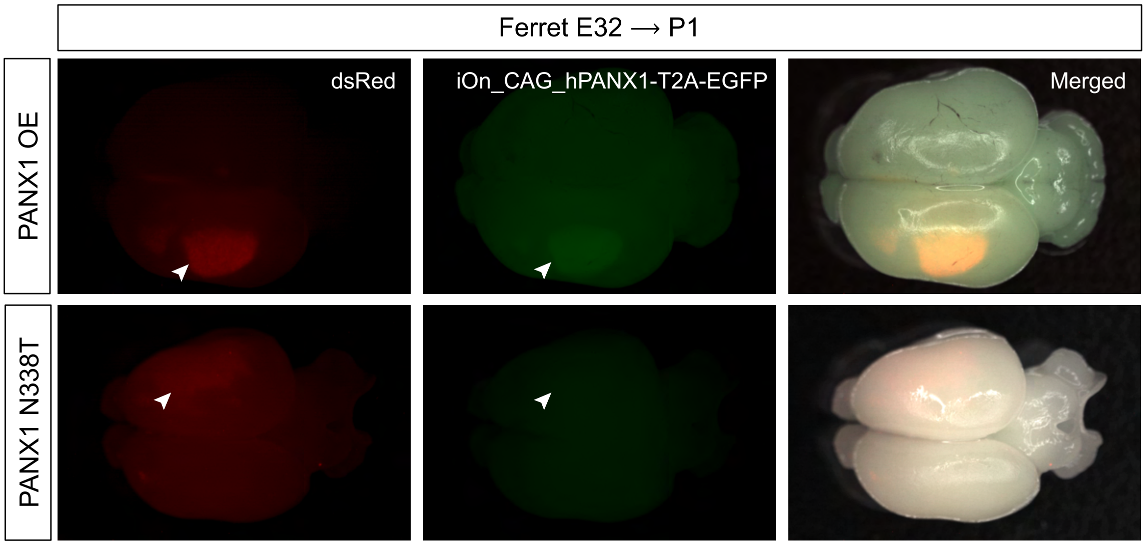


**Figure S12: *In utero* electroporation of mutant *PANX1* into ferret somatosensory cortex causes cell death.** Ferret embryos were electroporated at E32 with either WT or MT PANX1, along with pCAG-dsRed. Pregnant jills delivered naturally and kits were harvested on P1. *Top*, representative fluorescence imaging of P1 ferret cortex electroporated with WT PANX1 demonstrating overlap of dsRed and EGFP signal in the somatosensory cortex of the developing ferret brain (n = 8, WT). *Bottom*, representative fluorescence imaging of P1 ferret cortex electroporated with mutant PANX1 demonstrating a relatively weaker dsRed signal, and absent EGFP signal (n = 6, Asn338Thr). All 6 mutant-electroporated kits were identified based on dsRed signal and had no visible EGFP signal.

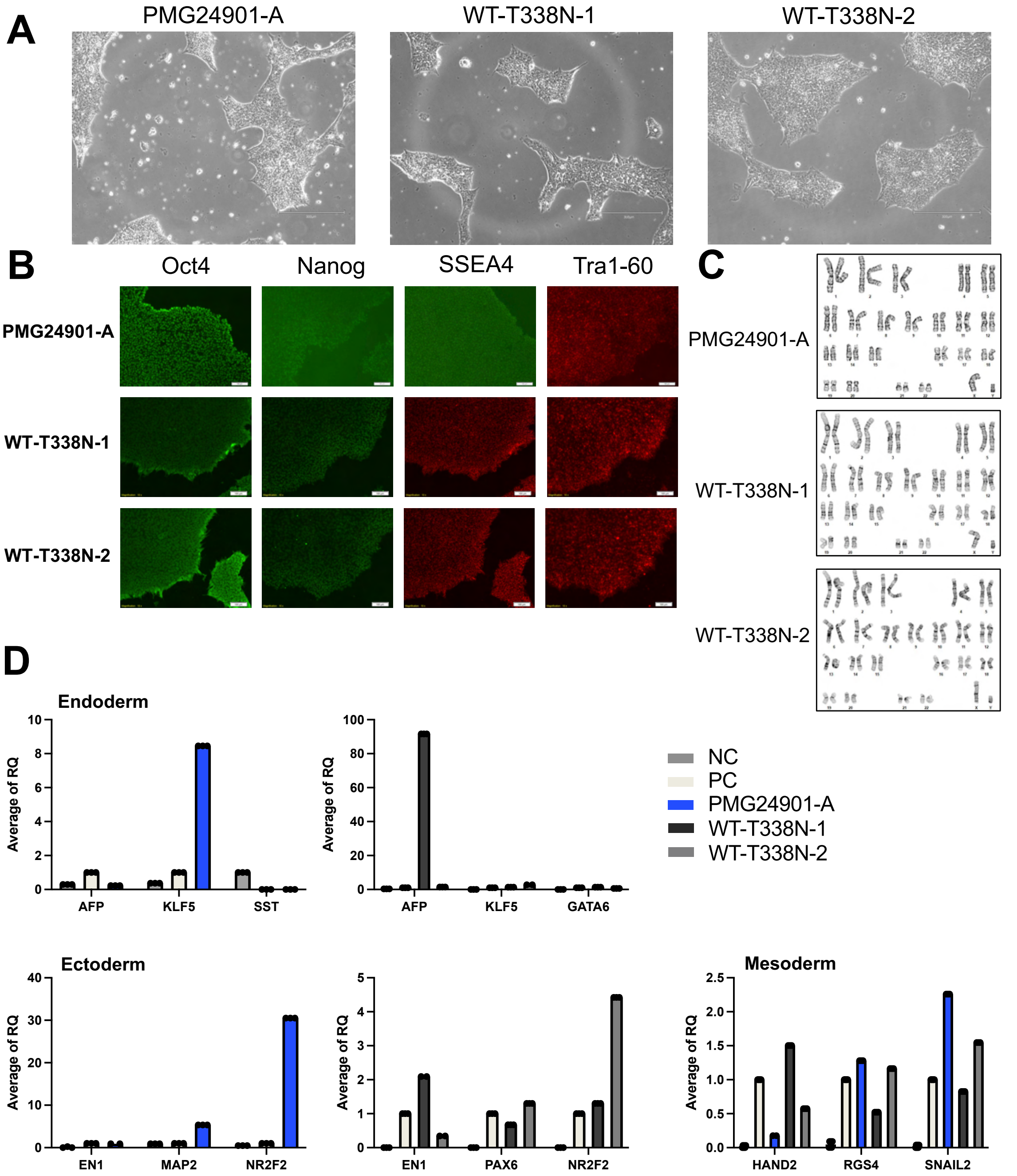


**Figure S13: Characterization of proband-derived iPSCs. (A)** Representative images of characteristic iPSC morphology two days after thawing cells in culture. Scale bar = 300μm. **(B)** Expression of pluripotency markers in PMG24901-A, WT-T338N-1, and WT-T338N-2. Scale bar = 100μm. **(C)** Normal 46XY karyotypes for each cell line. No clonal abnormalities detected. **(D)** qRT-PCR demonstrating expression of trilineage differentiation markers for each cell line. NC = negative control, PC = positive control. For endoderm and ectoderm, markers for WT lines placed on separate graphs as they were run with separate controls and marker sets.

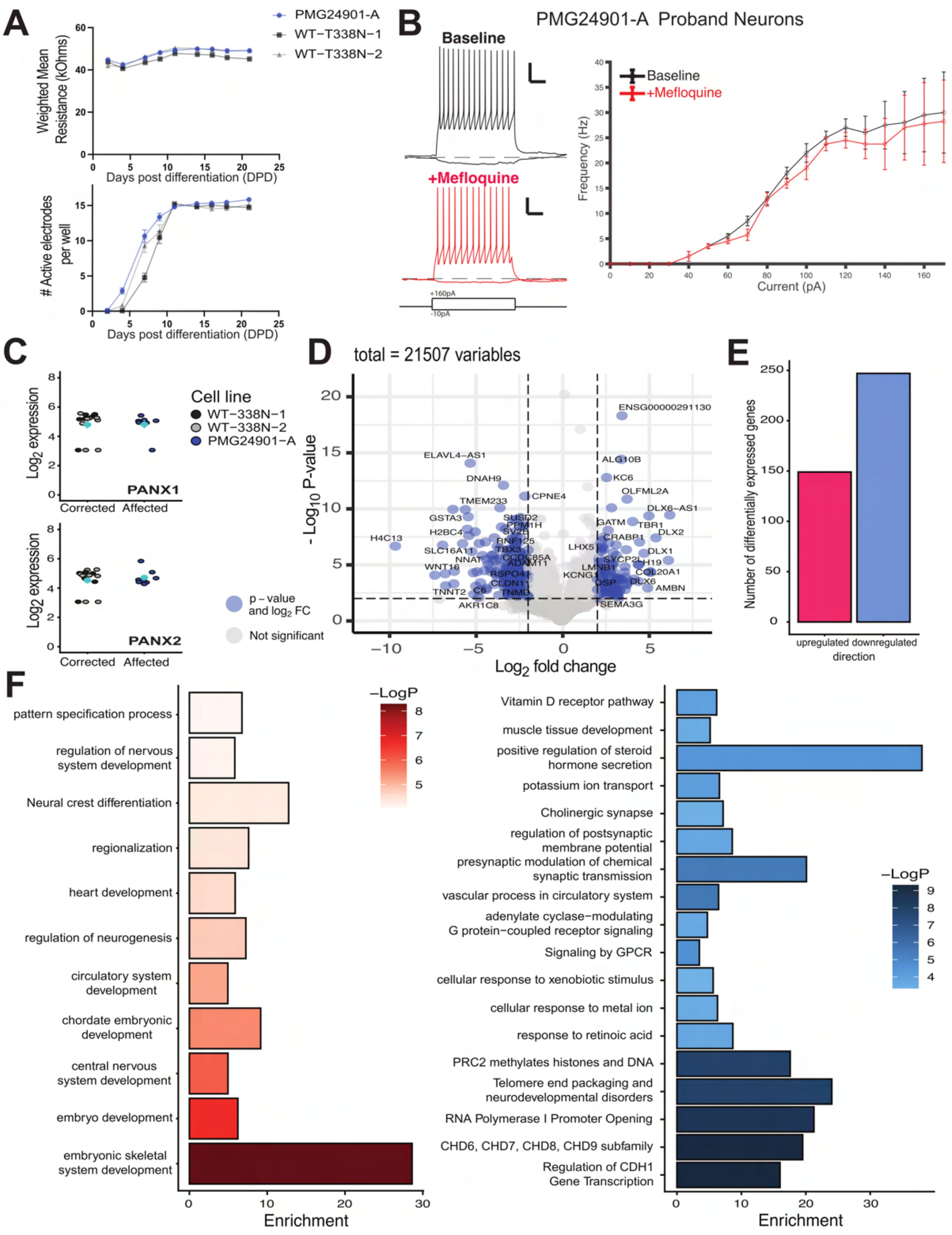


**Figure S14: Multielectrode array (MEA) recordings and bulk RNA-sequencing of iPSC-derived NGN2 neurons—related to Figure 4. (A)** *Top,* Consistent weighted mean resistance from a total of 16 wells in each of three replicate batches of NGN2 transduced neurons. *Bottom*, Number of active electrodes (out of 16 total) in each well for each cell line. Wells with fewer than 11 active electrodes by DPD 10 were excluded from the analysis. **(B)** *Left*, representative current clamp traces showing spiking from a proband neuron before and after mefloquine (1uM, 6 min)

perfusion, and (*Right*) summary analysis showing no change in the excitability curve (n=5, p >0.9). Scale bars 20 mv 100 ms. **(C)** Normalized log2 expression of PANX1 (upper panel) and PANX2 (lower panel) in proband-derived and two isogenic controls NGN2-induced neurons. Turquoise dots depict average expression of the corrected (n = 14) and affected (n=8) samples. **(D)** Volcano plot of differential expression analysis in DESEq2 depicting all tested genes. Blue circles correspond to the genes that passed both thresholds for significance (p < 0.01 and fold change |log2|> 2). **(E)** Bar plot indicating the total number of dysregulated genes in the affected genotype, divided by direction of change. **(F)** Functional enrichment analysis performed by querying multiple gene annotation databases, including Gene Ontology, Kegg Pathways, Reactome and Wikki pathways. Enrichment in the upregulated genes are shown in red (left panel) while the enrichment for the downregulated genes is shown in blue (right panel). Color scale indicates the significance level as the -Log10 p-value.

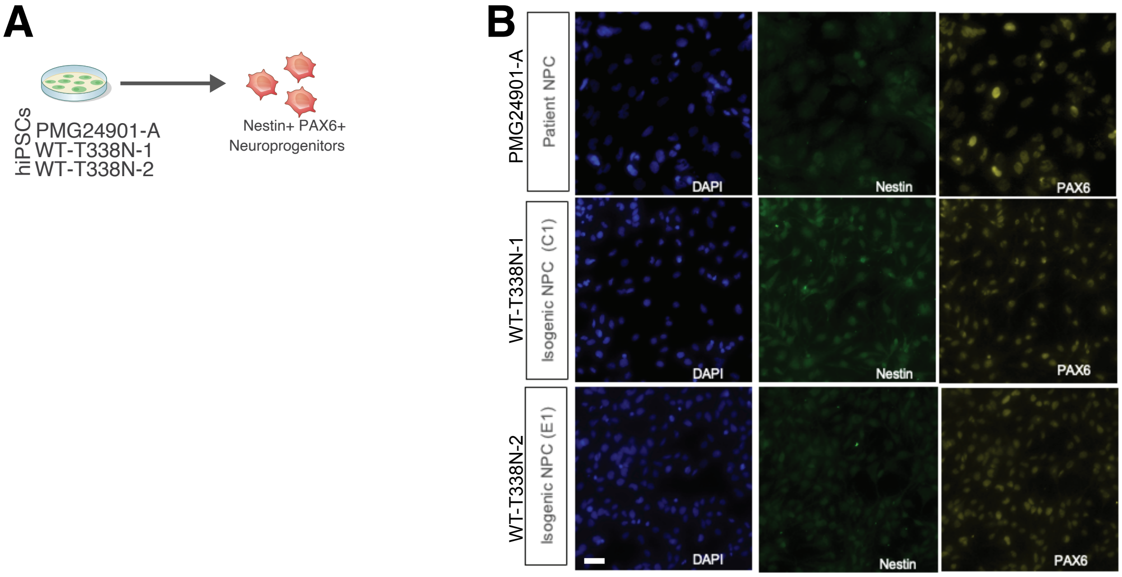


**Figure S15: Generation of NPCs from proband and isogenic controls**
**(A)** Schematic of NPC culture and immunocytochemistry. Proband (PMG24901-A) and isogenic WT (WT-T338N-1 and WT-T338N-2) cells were stained with Nestin and PAX6 for validation. **(B)** Representative imaging of NPCs from each genotype expressing Nestin and PAX6.

**Tables**

**Table S1.** Clinical case summaries of individuals with *PANX1* variants and PMG

| Individual ID | PMG20601 | PMGSL101 | PMG24901 |
|---|---|---|---|
| **Gene** | ***PANX1*** | ***PANX1*** | ***PANX1*** |
| **Genomic position (hg38)** | 11:94129352G>C | 11:94129422T>G | 11:94180069A>C |
| **cDNA change** | c.40G>C (NM_015368.4) | c.110T>G (NM_015368.4) | c.1013A>C (NM_015368.4) |
| **Protein change** | p.(Asp14His) | p.(Met37Arg) | p.(Asn338Thr) |
| **CADD v1.7** | 32 | 28.6 | 25.9 |
| **REVEL** | 0.656 | 0.284 | 0.607 |
| **PolyPhen-2** | 0.997 | 0.918 | 1 |
| **AlphaMissense** | 0.921 | 0.9011 | 0.9448 |
| **Mode of variant discovery** | Trio Exome | Trio Exome | Trio Exome |
| **Inheritance/Segregation** | Heterozygous, *de novo* | Heterozygous, *de novo* | Heterozygous, *de novo* |
| **Variant Classification (ACMG criteria)** | Likely Pathogenic (PS2, PM2, PP3) | Variant of Uncertain Significance (PM2, PM6, PP3) | Likely Pathogenic (PS2, PM2, PP3) |
| **Biological Sex** | Female | Female | Male |
| **Race/Ethnicity** | White/Not Hispanic of Latino | Hispanic | White/Not Hispanic of Latino |
| **Countries of Origin** | USA | El Salvador, Brazil | USA |
| **Parental consanguinity** | No | No | No |
| **Pregnancy history** | Maternal viral meningitis at 30-34wga, intrauterine growth retardation noted at 32wga | Uncomplicated, exposures denied | Uncomplicated |
| **Birth and neonatal history** | Spontaneous vaginal delivery, neonatal respiratory distress, 12d in NICU for apnea | Spontaneous vaginal delivery, healthy neonatal course, discharged home on day 2 | Spontaneous vaginal delivery, shoulder dystocia without fractures, 5 days in NICU after a cyanotic spell |
| **Estimated gestational age at delivery (weeks)** | 38 | 40 | 36 |
| **Birth weight** | 2.41 kg (-1.95 SD) | 3.204 kg (-0.25 SD) | 2.92 kg (67th percentile for 36 weeks gestation) |
| **Birth length** | 45.72 cm (-1.67 SD) | 49.5 cm (+0.10 SD) | NA |

| | | | |
|---|---|---|---|
| **Birth head circumference** | 30.5 cm (-2.92 SD) | 33 cm (-0.90 SD) | 31.75 cm (27th percentile for 36 weeks gestation) |
| **Age at last evaluation (#y#m)** | 1y1m | 6y7m | 5y |
| **Head Circumference** | 5m: 37 cm (-3.82 SD)<br>1y1m: 40 cm (-4.21 SD) | 6y7m: 48.4 cm (-1.94 SD) | 11m: 40.8 cm (-4.2 SD)<br>2y: 43 cm (-3.8 SD) |
| **Weight** | 1y1m: 9.36 kg (-0.42 SD) | 6y7m: 18.3 kg (-1.08 SD) | 11m: normal by report |
| **Height** | 1y1m: 73.15 cm (-0.54 SD) | 6y7m: 108.5 cm (-1.88 SD) | 11m: normal by report |
| **Developmental delay** | Yes, Global developmental delay | Yes, Global developmental delay | Yes, Global developmental delay<br>4m: rolled over; 8m: began to crawl; 11m: right pincer grasp; 1y: army crawl, sat with support, say mama and dada, play peek-a-boo; 19m: pull to stand and cruise, help with dressing; 2y: not yet walking independently, 5-10 words and 2 signs, used hands (right greater than left) to scribble, throw and use utensils to eat, point to body parts and to diaper when needed changing; 5y: attends school, running and jumping independently, communicates with signs and >20 single words |
| **Neuromuscular abnormalities** | Hypotonia | Central hypotonia, spastic quadriplegia cerebral palsy, hypertonia, decreased extension with tight hamstrings bilaterally, bilateral tapering of legs, poor head control | 6m: right arm preference with clenching of the left hand and decreased use of the left arm<br>2y: left-sided hemiparesis |

| | | | |
|---|---|---|---|
| | | | (upper extremity greater than lower) |
| **Epilepsy/Seizures** | Yes, Infantile spasms onset 3m, partial myoclonic seizures | None | Yes |
| **EEG abnormalities** | Yes, 3m: Grade III dysrhythmia with paroxysmal episodes that appear to be associated with desynchronization of the background, consistent with an underlying epileptiform disorder, but probably from a subcortical generator. In addition, there is a right central sharp wave which was continuous. 7m: Grade III dysrhythmia, the background is less disorganized than 2 weeks ago with a pattern approaching hypsarrhythmia. The universal high amplitudes from 2 weeks ago are no longer seen and the only higher amplitudes are slow waves out of left hemisphere, maximal activity for spike activity remains on left hemisphere but discontinuity in sleep has | Yes, 3y10m: Mildly abnormal EEG with slowing intermittently in centro-parietal location independently. The clinical concern of jerky movement was noted on this EEG, did not have any electrographic epileptiform discharges. This is not suggestive of epileptic spasm or myoclonic jerk of central origin based on this EEG. | Yes, 3y7m: consistent with epileptiform activity and generalized and multifocal epilepsy, 4y3m: tonic seizures recorded during sleep |

| | | | |
|---|---|---|---|
| | reduced. This study, although abnormal, is beginning to approach the abnormalities identified on study at 3m. | | |
| **Dysmorphic features** | | 6y7m: microcephalic, sloping forehead, inter-canthal distance measure 3.4 cm, left palpebral fissure measured 2.8 cm, right palpebral fissure measured 2.9 cm, left ear measured 5.7 cm, right ear measured 6.3 cm, prominent lateral palatine ridges, left hand with hockey stick palmar crease | |
| **Eye/vision or Ear/hearing abnormalities** | Cortical visual impairment | Stable bilateral hyperopia, (normal hearing) | Myopia and right esotropia, wears glasses |
| **Cardiovascular abnormalities** | | None | |
| **Gastrointestinal and Genitourinary abnormalities** | | Gastrostomy tube | |
| **Skeletal and Connective tissue abnormalities** | | Pathologic dislocation of right hip joint and scoliosis (27 degree left upper thoracic curve and 21 degree right lower thoracic curve) | 20m: x-ray questioned diminished thoracic kyphosis and lumbar lordosis (normal lumbar spine and pelvis) |
| **Other diangoses or findings** | | Obstructive sleep apnea | 11m: left arm and leg reportedly smaller in size than on right; 2y: drooling, bilateral ankle foot orthoses |

| | | | |
|---|---|---|---|
| **Brain imaging findings** | MRI at 12m and 26m: extensive, bilateral polymicrogyria centered in the perisylvian region and extending over the convexities, more significantly on the right than the left. Severely depressed white matter volume, moderately depressed thalamic volume, and mild thinning of the brainstem are also noted. Severe callosal agenesis with moderate-severe reduction in length and immeasurably thin. Mild inferior vermian hypoplasia was present. Myelination was age-appropriate though there was minimal periventricular white matter dysmyelination versus prominence of the perivascular spaces. Basal ganglia were normally cleaved though borderline depressed in volume. Findings are the same at 12 and 26 months except myelination had | MRI at 5m: extensive bilateral perisylvian polymicrogyria extending over frontal and temporal lobes. White matter volume was moderately depleted, and thalamic volume was somewhat reduced. Frontal lobe white matter was more T2 hyperintense than expected for age-expected incomplete myelination, raising the possibility of dysmyelination or gliosis. The basal ganglia, pituitary (allowing for an incidental pars intermedia cyst), brainstem, cerebellum, and orbital and olfactory structures were within normal limits. | MRI at 10m: small right hemisphere with diffuse polymicrogyria, reduced white matter and thalamic volume, and prominent perivascular spaces (some dysmyelination not excluded). Basal ganglia, corpus callosum, cerebellum and optic nerves within normal limits. The presence or absence of a left-sided cortical malformation was difficult to appreciate, however repeat imaging at 4 years 5 months was reported to reveal polymicrogyria involving the parasagittal left frontal lobe, from the left orbital frontal parenchyma posteriorly to the left paracentral lobule, and partially involving the left cingulate gyrus (this 10 month MRI with thick slices suggests polymicrogyria at the depth of a left frontal sulcus and the posterior left sylvian fissure though it is difficult to be certain). |

| | | | |
|---|---|---|---|
| | matured on the older study. | | |
| **Prior genetic/laboratory testing** | Karyotype, 46,XX, Chromosome microarray (2012): gain of 15q13.2-q13.3(28, 680, 834-30, 340, 762) overlapping CHRNA7. Clinical WES (2014): Non-diagnostic variants noted only | Chromosome microarray (2019): arr[GRCh37] (1-22,X)x2 Normal Female | Normal newborn metabolic screening, basic metabolic panel, thyroid function and ceruloplasmin, chromosome microarray, cortical brain malformation gene sequencing panel (with deletion/duplication analysis) |

**NA:** Not available or not assessed

**Table S2.** List of all reported variants in *PANX1*.

| Disease Association | PANX1 Variant | Reference | Inheritance |
|---|---|---|---|
| Infertility | 21_23delTEP | Sang et al. (5) | Unknown |
| Infertility | Arg29Gln | Wu et al. (7) | AD |
| Infertility | Ser137Leu | Zhou et al. (9) | AD |
| Infertility | Ser238Pro | Wang et al. (6) | AR |
| Infertility | Arg300Gln | Wang et al. (6) | AR |
| Infertility | Asn326del | Zhou et al. (8) | AD |
| Infertility | Lys346Glu | Sang et al. (5) | AD |
| Infertility | Cys347Ser | Sang et al. (5) | AD |
| Infertility | Gln392* | Sang et al. (5) | AD |
| PMG | Asp14His | Akula et al. (27) | *De novo* |
| PMG | Met37Arg | Akula et al. (27) | *De novo* |
| PMG | Asn338Thr | Akula et al. (27) | *De novo* |
| VUS | Arg217His | Shao et al. (11) | AR |

**AD**, autosomal dominant; **AR**, autosomal recessive

**Supplementary Dataset 1 (separate file).** Averaged amplitudes and spike counts per well during a representative 10 min multielectrode array recording.

**Supplementary Dataset 2 (separate file).** Dataset from bulk RNA-sequencing of NGN2-derived neurons: differentially expressed genes, enrichment analysis, and sequencing statistics

**Movie S1 (separate file)—Related to Figure 2.** Time-lapse confocal imaging of migrating EGFP+ and dsRed+ cells in an E16.5 (E14.5 IUE) embryonic mouse cortex electroporated with control constructs.

**Movie S2 (separate file)—Related to Figure 2.** Time-lapse confocal imaging of migrating EGFP+ and dsRed+ cells in an E16.5 (E14.5 IUE) embryonic mouse cortex electroporated with WT PANX1 and dsRed.

**Movie S3 (separate file)—Related to Figure 2.** Time-lapse confocal imaging of migrating EGFP+ and dsRed+ cells in an E16.5 (E14.5 IUE) embryonic mouse cortex electroporated with N338T PANX1 and dsRed.

**Supplementary Methods**

**KEY RESOURCES TABLE**

| REAGENT or RESOURCE | SOURCE | IDENTIFIER |
|---|---|---|
| **Antibodies** | | |
| Rabbit Polyclonal PAX6 | ThermoFisher | Cat#42-6600; RRID: AB_2533534 |
| Rabbit Monoclonal CC3 | Cell Signaling Technology | Cat#9664S; RRID: AB_2070042 |
| Mouse Monoclonal Ki-67 | BD Biosciences | Cat#550609; RRID: AB_393778 |
| Rat Monoclonal Ctip2 | Abcam | Cat#ab18465; RRID: AB_2064130 |
| Mouse Monoclonal SATB2 | Abcam | Cat#ab51502; RRID: AB_882455 |
| Chicken Polyclonal GFP | Aves Labs | Cat#GFP-1020; RRID: AB_2307313 |
| Rabbit Polyclonal RFP | Rockland | Cat#600-401-379; RRID: AB_2209751 |
| Goat Polyclonal SOX2 | R&D Systems | Cat#AF2018; RRID: AB_355110 |
| Guinea Pig Monoclonal c-Fos | Synaptic Systems | Cat#226 308; RRID: AB_2905595 |
| Mouse Monoclonal RFP | ThermoFisher | Cat#MA5-15257; RRID: AB_10999796 |
| Goat Polyclonal RFP | Rockland | Cat#200-101-379; RRID: AB_2744552 |
| Rabbit Monoclonal PANX1 | Cell Signaling Technology | Cat#91137S; RRID: AB_2800167 |
| Rabbit Polyclonal PANX1 | Sigma-Aldrich | Cat#HPA016930; RRID: AB_1854954 |
| Mouse Monoclonal ROR-beta | R&D Systems | Cat#PP-N7927-00; RRID: AB_3659610 |
| Rabbit Monoclonal Brn2/POU3F2 | Cell Signaling Technology | Cat#12137S; RRID: AB_2797827 |
| Mouse Monoclonal Vinculin | Sigma Aldrich | Cat#V913; RRID: AB_477629 |
| Mouse Polyclonal MAP2 | Sigma Aldrich | Cat#M9942; RRID: AB_477256 |
| Chicken Polyclonal MAP2 | Invitrogen | Cat#PA1-10005; RRID: AB_1076848 |
| **Bacterial and viral strains** | | |
| TetO-Ngn2-Puro | Addgene | #52047 |

| | | |
|---|---|---|
| FUW-M2rtTA | Addgene | #20342 |
| **Chemicals, peptides, and recombinant proteins** | | |
| DAPI solution | ThermoFisher | 62248 |
| ProLong Diamond Antifade Mountant | ThermoFisher | P36961 |
| Carbenoxolone disodium salt | Sigma-Aldrich | C4790 |
| 10Panx | Tocris Bioscience | 3348 |
| Fast Green FCF | Sigma-Aldrich | F7252 |
| Lipofectamine 3000 Transfection Reagent | ThermoFisher | L30000015 |
| ARL 67156 trisodium salt hydrate | Sigma-Aldrich | A265 |
| BDNF | Peprotech | Cat#450-02 |
| NT3 | Peprotech | Cat#450-03 |
| Laminin | Life Technologies | Cat#23017-015 |
| Ara-C | Sigma-Aldrich | Cat#C1768 |
| BzATP triethylammonium salt | Tocris | Cat#33-121 |
| YO-PRO-1 Iodide | Invitrogen | Cat#Y3603 |
| Trizol Reagent | Invitrogen | Cat#15596018 |
| 1-Bromo-3-chloropropane | Sigma-Aldrich | Cat#B9673-200ML |
| EDTA (0.5M), pH 8.0, RNAse-free | Invitrogen | Cat#AM9260G |
| 2-Propanol, Molecular Biology Grade | Fisher BioReagents | Cat#BP2618500 |
| Ethanol, Absolute (200 Proof), Molecular Biology Grade | Fisher BioReagents | Cat#BP2818500 |
| Nuclease Free Water | Invitrogen | Cat#AM9939 |

| | | |
|---|---|---|
| HBSS, no calcium, no magnesium, no phenol red | Gibco | Cat#14-175-095 |
| HEPES (1M) | Gibco | Cat#15-630-080 |
| Mefloqune Hydrochloride | Tocris | Cat#6819 |
| **Critical commercial assays** | | |
| Pierce™ BCA Protein Assay Kit | ThermoFisher | Cat#23227 |
| ATP Determination Kit | Invitrogen | Cat#A22066 |
| Click-iT™ Plus EdU Alexa Fluor™ 647 Imaging Kit | Invitrogen | Cat#C10640 |
| Jess Automated Western Blot 12-230kDa (Chemiluminescent) | Bio-Techne | Cat#SM-W003 |
| **Deposited data** | | |
| Bulk gene expression of human developing cortex | Allen Brain Atlas | |
| Single cell atlas of developing human cortex | Wang et al. (23) | nemo:dat-mbkuyiy |
| Single cell atlas of ferret cortex | Bilgic et al. (24) | DDBJ: DRA016867 |
| **Experimental models: Organisms/strains** | | |
| Mouse: CD-1 | Charles River | 022CD1 |
| Ferret: domesticated ferret | Marshall Biosciences | SPF Ferrets |
| **Oligonucleotides** | | |
| RNAscope Probe-Hs-PANX1 | ACD Biotechne | #589911 |
| RNAscope Probe-Hs-GFAP | ACD Biotechne | #311801 |
| **Recombinant DNA** | | |
| iOn_CAG_hPANX1 WT and variants | This study | N/A |
| piggyBase | Yusa et al. (26) | Gift from the Cepko Lab |
| pCAG-dsRed | This study | N/A |
| **Software and algorithms** | | |

| | | |
|---|---|---|
| GraphPad | Prism | https://www.graphpad.com/ |
| Fiji | ImageJ | https://imagej.net/software/fiji/ |
| Zen | Zeiss | https://www.zeiss.com/microscopy/us/products/software/zeiss-zen.html |
| MATLAB | MathWorks | https://www.mathworks.com/products/matlab.html |
| Python | Python | https://www.python.org/ |
| R | R-project | https://www.r-project.org/ |
| bioRender | bioRender | https://www.biorender.com/ |
| Illustrator | Adobe | https://www.adobe.com/ |
| Metascape | Metascape | https://metascape.org/gp/index.html#/main/step1 |
| Sciugo | Sciugo | https://sciugo.com/ |
| Zeiss Arivis Pro/ Vision4D | Zeiss | https://www.zeiss.com/microscopy/us/products/software/arivis-pro.html |
| Nikon NIS-Elements | Nikon | https://www.microscope.healthcare.nikon.com/products/software/nis-elements |
| Axion Integrated Studio | Axion Biosystems | https://www.axionbiosystems.com/resources/product-brochure/axis-brochure |
| MetaXpress | Molecular Devices | https://www.moleculardevices.com/ |
| Incucyte Live Cell Imaging and Analysis | Sartorius | https://www.sartorius.com/en/products/live-cell-imaging-analysis/live-cell-analysis-software |

**Resource availability**

Further information and requests for resources and reagents should be directed to and will be fulfilled by Christopher A. Walsh or Richard S. Smith. Plasmids generated in this study are available from the lead contact with a completed materials transfer agreement. Accession numbers for previously published RNA-seq datasets are listed in the Key Resources Table. Original western blot images and microscopy data have been deposited at Mendeley (DOI: 10.17632/yzn5523wvn.2). This paper does not report original code.

**Animal husbandry**

Timed pregnancies of CD1 mice were obtained from Charles River Laboratories and analyzed from embryonic through early postnatal ages; mice of both sexes were used. Ferrets were obtained from Marshall Bioresources and same-sex housed with a dedicated Large Animal Facility on a 12-h light/dark cycle at 18-23 °C with rotating sensory enrichment, and provided food and water ad libitum.

**BrainSpan, RNAscope, MERFISH, and slide immunostaining**

For BrainSpan analyses, *PANX1* (chr11:94128841-94181968, GRCh38/hg38), *PANX2* (chr22:50170731-50180295, GRCh38/hg38), and *PANX3* (chr11:124611428-124620356, GRCh38/hg38) RPKM (reads per kilobase exon per million mapped reads) values were obtained for annotated neocortical tissues spanning 8 pcw to 19 y and fit with a polynomial in R. Brain regions included dorsolateral prefrontal cortex; ventrolateral prefrontal cortex; anterior (rostral) cingulate (medial prefrontal) cortex; orbital frontal cortex; primary motor-sensory cortex; parietal neocortex; posterior (caudal) superior temporal cortex (area 22c), inferolateral temporal cortex (area 20); occipital neocortex, thalamic regions and hippocampus, among others.

For RNAscope, human fetal brains (17-20 gestational weeks, both sexes) were fixed in 4% PFA, cryoprotected in 30% sucrose, frozen in isopentane, and sectioned at 20-30 µm (Leica Cryostat) and mounted onto warm charged SuperFrost Plus slides (Fisher). Multiplex fluorescent RNAscope was performed using the manufacturer’s protocol (ACD Multiplex v2) with PANX1 (589911) and GFAP (311801) probes. Images were acquired at 20x on a Zeiss Axio Observer, tiled, stitched, and analyzed in Zen Blue.

For detailed descriptions of human tissue preparation, annotation, and MERFISH gene-panel selection, imaging, and data processing, see Ref. (22). Briefly, 10-µm de-identified fresh-frozen human fetal tissue sections were hybridized with a 960-gene Vizgen panel for 36 h at 37 °C, gel embedded, proteinase-K cleared, stained with DAPI/PolyT, and imaged according to the MERSCOPE Instrument Preparation guide. Nuclei were segmented before transcript filtering and normalization.

For slide immunostaining, fresh-frozen tissue was washed with PBS and fixed on-slide in 4% PFA for 15 min at RT, subjected to heat-mediated antigen retrieval, and blocked in 8% normal donkey serum + 0.3% BSA and 0.3% Triton X-100 for 1 hour. Primary antibodies were incubated overnight at 4 °C and secondary antibodies conjugated to Alexa fluorophores + DAPI for 2 h at room temperature. Confocal imaging used a Zeiss LSM 980 or Axioscan 7 Slide Scanner; 20x tiled images were stitched and analyzed using Zen Blue and Fiji.

**Single-cell RNA-seq analysis**

The developmental neocortex single-nucleus multiome gene expression dataset (23) was analyzed using the supplied developmental-stage, region, and cell-type annotations. For targeted *PANX1* analysis, the RNA assay was log-normalized, and expression was quantified across cell types as both mean normalized expression and the percentage of PANX1-positive cells. Feature plots, dot plots, fetal/postnatal and PFC/V1 comparisons, and developmental line plots were generated in Seurat, dplyr, and ggplot2.

Single-cell RNA-seq atlas of ferret cortex was obtained from Ref. (24) and similarly analyzed for targeted *PANX1* analysis using the supplied developmental-stage and cell-type annotations.

**Molecular cloning and cell culture**

*PANX1* constructs were constructed by VectorBuilder; hyperactive piggyBac transposase was donated as a gift by the Cepko laboratory. Esp3I-T2A-EGFP was cloned into iOn-CAG∞MCS (Addgene 154013) between transposase recognition sites using BamHI/AvrII. *PANX1* was PCR-amplified with Esp3I sites (5'-ttaaCGTCTCcgtacgccaccatggccatcgctca; 3'-ttaaCGTCTCCCCTCgcaagaagaatccagaagtctctgt), inserted into iOn-CAG-Esp3I-T2A-EGFP, and sequence verified. Mutant constructs were generated by GenScript bridge-PCR mutagenesis and validated with full-length plasmid sequencing.

N2A (ATCC, CCL-131) and HEK293T (ATCC, CRL-3216) cells were maintained at 37 °C and 5% CO2 in DMEM with 10% FBS and 1X penicillin/streptomycin. Cells were transfected with WT or mutant *PANX1* as well as piggyBase at a 5:1 ratio using Lipofectamine 3000; HEK293T cells used for electrophysiology were electroporated with the Neon system at a 4:1 *PANX1*:piggyBase ratio.

**Western blotting, BCA, and ATP determination**

HEK293T cells were harvested 48 h post transfection in M-PER (Thermo Fisher Scientific), separated on 4-20% Mini-PROTEAN TGX gels (Bio-Rad Cat. 4568094), and transferred to PVDF membranes. Membranes were blocked with LI-COR Intercept buffer (Cat. 927-60001) and probed with rabbit anti-PANX1 (Sigma HPA016930, 1:200), mouse anti-vinculin (Sigma V9131, 1:500), and chicken anti-GFP (Aves GFP-1020, 1:500), followed by appropriate IRDye secondaries. Blots were imaged on a LI-COR Odyssey and quantified in Sciugo; PANX1 signal was normalized to GFP. For BCA assays, M-PER lysates were clarified at 12,000 rpm for 15 min at 4 °C and assayed with the Pierce BCA kit according to the manufacturer.

For ATP assays, transfected N2A cells underwent fluorescence-activated cell sorting (FACS) to isolate GFP-positive fractions and replated. Cells were placed in 2% heat-inactivated FBS at least 24 h before analysis, incubated with 100 µM ARL67156 (Sigma Cat. A265) for 20 min, and pretreated with 100 µM carbenoxolone (CBX; Sigma Cat. C4790) where indicated. ATP standards (0-100 nM) and 10-µL samples were assayed with 100 µL ATP reaction solution for 10-15 min in the dark before luminescence measurement on a SpectraMax iD5 (Molecular Devices). ATP release was normalized to total protein.

**Electrophysiology recordings of HEK293T cells**

Whole cell voltage clamp recordings were performed using Sutter Instruments IPA. Glass pipettes were forged from borosilicate glass on a Sutter Puller (P-2000) and filled with intracellular recording solution (in mM: 80 CsMES, 25 NaCl, 10 HEPES, 10 Cs4-BAPTA, 2 MgCl2, free Ca 90uM, OsM 295 mOs. pH 7.3 with CsOH). Pipettes were measured to have resistances of 3-4

MΩ prior to seal formation. Extracellular solution contained (mM): 150 NaCl, 10 HEPES, 1.8 $CaCl_2$; 300 mOsm, pH 7.4. Membrane resistance was derived from steady-state voltage responses to -10, -20, and -30 pA current injections; capacitance was calculated from the fitted membrane time constant ($\tau$ = RC). Voltage ramps extended from -90 to +100 mV over 600 ms and IV curves were generated in MATLAB.

**Mouse IUE, tissue preparation, and slice imaging**

At E14.5, approximately 2 µL of transposon plasmid (2.5 µg/µL), piggyBase (500 ng/µL), pCAG-dsRed (1 µg/µL), and 0.1% FastGreen FCF (Sigma, Cat. F7252-5G) was injected into the lateral ventricle followed by 5 x 50-ms, 50-V pulses at 1.1-s intervals (28) using paddle electrodes oriented over the dorsal surface of skull. Pregnant dams were euthanized with $CO_2$ and embryos were harvested at embryonic day 17.5 (E17.5). E17.5 brains were drop-fixed overnight in 4% PFA; P10 animals were anesthetized with ketamine and xylazine, transcardially perfused with PBS and 4% PFA, and brains fixed for 2-5 h. Tissue was cryoprotected sequentially in 15% and 30% sucrose, embedded in OCT, sectioned at 18 µm, and mounted on charged slides.

For time-lapse imaging experiments, embryos were collected 48 h after E14.5 IUE. Fluorescent brains were embedded in 3% low-melting-point agarose, sectioned at 300 µm on a vibratome (Leica VT1000S), and cultured on 0.4-µm transwells (Millicell PICM03050) in a glass bottom cell culture dish (FluoroDish 5040-10) in DMEM/F12 without phenol red and supplemented with 10% FBS, penicillin/streptomycin, and B27. Multipoint z-stacks were acquired every 10 min for up to 24 h on a Dragonfly Spinning Disk confocal microscope and images stitched and analyzed in Imaris.

**Ferret postnatal and in utero electroporation**

For P1-P2 postnatal electroporation (29), kits were deeply anesthetized with isoflurane, bupivacaine administered subcutaneously into the scalp, and positioned stereotaxically (Stoelting, Cat. #51615). Under sterile surgical conditions, the skin over the head was cut along the midline, and a small opening was made on the skull (-0.5 mm posterior and +2.0 mm lateral to the

Bregma) using a micro knife (Ambler Surgical, Cat. #964501). ~3 µL plasmid mixture (WT or mutant transposon 2.5 µg/µL, pCAG-PBase 500 ng/µL, pCAG-dsRed 1 µg/µL, 0.1% FastGreen) was injected 2 mm below pia into the right lateral ventricle. Tweezers with needle and paddle electrodes (NEPA Gene, Cat. #CUY661-3X7) were used to electroporate, delivering five 50-ms, 50-V pulses at 1-s intervals. The injection micropipette and the electrode tweezers were each held with two separate micromanipulator arms assembled into the stereotaxic frame (Stoelting, Cat. #51606 and #51631 for the micropipette, Cat. #51604 and #51634 for the electrode). The micropipette holder and electrode holder were angled 22.5° to target prospective occipital cortex. Kits were recovered, monitored, and returned to the dam.

Ferret IUE was performed at E32-E33 according to standard procedures (30). Briefly, pregnant jills were anesthetized with isoflurane, embryos exposed by laparotomy, and ~3 µL of the same plasmid mixture injected intraventricularly. Five 100-V square pulses were delivered at 1-s intervals. Tissue was harvested at P1 or P21, fixed in 4% PFA, and successful transfection assessed by GFP/RFP fluorescence. Both sexes were targeted.

**Ferret tissue preparation**

Ferret kits were deeply anesthetized with ketamine/xylazine and transcardially perfused with PBS followed by 4% PFA. Dissected brains were postfixed for 24 h and cryoprotected in 15-30% sucrose. P1 brains were cryosectioned at 20 µm and mounted immediately onto warm, charged SuperFrost Plus slides; older brains were cut at 50 µm on a sliding microtome (Leica SM2010 R) and stored overnight at -20ºC in anti-freezing solution. Free-floating sections were permeabilized with 0.25% Triton X-100, blocked in 8% normal donkey serum + 0.3% BSA + 0.3% Triton X-100, incubated with primary antibodies overnight at 4 °C and treated with secondary antibodies plus DAPI for 2 h, and imaged on a Zeiss LSM 980.

**iPSC generation, correction, and maintenance**

The PBMCs were thawed, and the erythroblast population was expanded for 9 days in Stem Span SFEM II medium with erythroid expansion supplement (Stem Cell Technologies, Cat #9605 and #2692). On day 9, 100,000 cells were transduced using the CytoTune™-iPS 2.0 Sendai Reprogramming Kit (Thermo Fisher) following the manufacturer instructions. Four days later, the transduced cells were transferred to a 10cm plate with irradiated MEFs in the following medium: DMEM/F12, 20% KO-SR, NEAA 0.1mM, L-Glutamine 2mM, Beta Mercaptoethanol 1x and 10ng/ml of bFGF. Colonies were picked after 3 wk, then expanded on Cultrex (Biotechne, Cat# 3434-005-02) and mTeSR+ medium (Stem Cell Technologies Cat# 05825). Pluripotency was verified by qRT-PCR (Oct4, Sox2, Nanog, hTERT, DnmT3B), immunostaining (Oct4, Nanog, SSEA4, TRA-1-60), embryoid-body tri-lineage differentiation, normal G-banded karyotype (Wicell), normal BCL2L1 copy number, and negative mycoplasma testing.

CRISPR-Cas9 correction of PANX1 Thr338Asn was performed by EditCo Bio (Redwood City, CA, USA). To generate these cells, Ribonucleoproteins containing the Cas9 protein and synthetic chemically modified guide RNA produced by Synthego were electroporated into the cells along with a single-stranded oligodeoxynucleotide (ssODN) donor using EditCo's optimized protocol. Editing was assessed after 48 h by PCR and Sanger sequencing or NGS; monoclonal cell populations were generated by single-cell deposition and verified using the PCR-Sanger-ICE genotyping strategy above. Feeder-free iPSCs were maintained in mTeSR Plus on Geltrex, passaged with Accutase, and monitored for mycoplasma and karyotype abnormalities. The iPSCs used throughout the study were below passage 50. All studies were performed with approved protocols of Boston Children's Hospital.

**NGN2 induction and neuronal maintenance**

hiPSCs were infected overnight with TetO-Ngn2-Puro (Addgene plasmid #52047) and FUW-M2rtTA (Addgene plasmid #20342) lentiviruses, produced by the Viral Core at Boston Children's Hospital. The viral mixture was added to the cells in mTeSR Plus medium, supplemented with 10 µM ROCK inhibitor Y-27632 (Cayman, catalog #10005583) and 8 µg/mL polybrene (Sigma-Aldrich, catalog #TR-1003-G), expanded for two passages, banked, and replated at 124,000 cells/cm2. On day 0, cells received N2 medium with 2 µg/mL doxycycline and BDNF/NT3; on day 1, 1 µg/mL puromycin was added. On days 2-3, cultures were switched to Neurobasal

A/B27/GlutaMAX containing doxycycline, puromycin, BDNF/NT3, and 2 µg/mL Ara-C. On day 6, cells were dissociated with papain/DNase I and replated for downstream assays.

For MEA studies, day-6 neurons were seeded with human astrocytes in 10 µL droplets containing 100,000 neurons/well, 15,000 astrocytes/well (NCardia, catalog #M0605), 20 µg/mL laminin (Thermofisher, catalog #23017-015), 10 µM ROCK inhibitor, and 1x CEPT cocktail (Tocris, catalog #7991). Neurons were maintained with astrocyte-conditioned medium prepared from NCardia astrocytes and diluted 1:1 with Neurobasal A + 1x GlutaMAX + 1x B27, supplemented with BDNF and NT3; feeds were performed every other day. Importantly, figures use days-post-differentiaion (DPD), which represents days after DOX induction.

**Neuronal immunocytochemistry and automated morphometry**

Day-6 neurons were plated on poly-ornithine/laminin-coated 96-well plates at 40,000 cells/well and fixed after 24 h in 4% PFA and washed with dPBS. Cells were blocked in 5% normal goat serum, 2% BSA, and 0.1% Triton X-100 and incubated overnight at 4°C with anti-MAP2 (Sigma M9942, 1:2000), followed by Alexa-fluorophore secondary antibodies and Hoechst 33258. Imaging was performed at 10× magnification without a z-stack capturing nine fields/well across eight wells/line were acquired at 10x on an ImageXpress MicroXLS and analyzed with the MetaXpress neurite algorithm.

**Patch clamp of NGN2 neurons**

Recordings were performed at DPD 26-32 using 3-5 MΩ borosilicate pipettes. Internal solution contained (mM): 97.5 K-gluconate, 32.5 KCl, 10 HEPES, 1 mM EGTA, 2 MgCl2, plus 1 mM Alexa Fluor 488; adjusted to 295 mOsm, pH 7.4 with KOH. The internal solution also contained Alexa-Fluor 488 nm dye to visualize neuron morphology (1 mM, Thermo Fisher). ACSF contained (mM): 125 NaCl, 2.5 KCl, 2 CaCl2, 1 MgCl2, 1.25 NaH2PO4, 26 NaHCO3, 15 glucose, 1 myo-inositol, 2 sodium pyruvate, and 0.4 ascorbic acid; 300 mOsm, pH 7.4, (adjusted with NaOH), continuously oxygenated with 95% O2/5% CO2.

### Multielectrode-array acquisition and analysis

CytoView 48-well plates (M768- tMEA-48B; Axion Biosystems) were coated with 0.1% PEI in borate buffer. Day-6 neurons were seeded at 110,000 neurons plus 16,500 astrocytes/well with laminin, Y-27632, and CEPT cocktail. Recordings of spontaneous network activity began 2 d after seeding and continued on M/W/F schedule for 25 d; half-media changes occurred 24 h before recording. Plates acclimated for 10 min at 37 °C/5% CO2 before each 10-min recording. Data were sampled and analyzed using Axion Integrated Studio (AxIS) at 12.5 kHz and filtered at 200-3,000 Hz; adaptive spike detection was set at 6 SD/electrode. Network-burst settings were 100-ms maximum interspike interval, >=50 spikes, and >=35% participating electrodes. Parameter trajectories were fit by nonlinear least squares to three-parameter sigmoidal curves (midpoint, slope, peak) and coefficients compared across genotypes in R.

Threshold-detected spikes were exported without spike sorting from Axis Navigator. Average raw spike amplitudes and spike counts during a representative 10-min recording are reported in Supplementary Dataset 1. Weighted mean firing rate excluded inactive electrodes. Multi-electrode simultaneity was calculated from cross-correlograms for all electrode pairs, pooled and normalized for spiking regularity using a 20-ms window.

### Bulk RNA-seq processing

Monolayer neuronal cultures were lysed in TRIzol, phase separated with 1-bromo-3-chloropropane, and RNA precipitated with 2-propanol, washed in 70% ethanol, and resuspended in nuclease-free water with 0.1 mM EDTA. Samples were sent to plasmidsaurus for bulk RNA sequencing. DESeq2 analyses modeled a negative binomial distribution; sequencing run and read depth were included as batch covariates because preliminary analyses indicated strong associated batch effects (Supplementary Dataset 2). Metascape enrichment used default parameters across Gene Ontology, Reactome, WikiPathways, and KEGG.

**NPC immunocytochemistry, western blot, and YO-PRO-1 uptake assay**

iPSC-derived NPCs were generated using STEMCELL Technologies neural induction and maintenance kits, following a 20-day differentiation protocol to yield stable Nestin+/PAX6+ neural progenitor cells (See Manufacturers protocol). For immunocytochemistry, cells were processed following the same protocol as iPSC-derived neurons and stained with anti-PAX6 (Invitrogen MA1109, 1:100) and anti-PANX1 (Cell Signaling Technology 91137, 1:100), followed by anti-rabbit Alexa 488 and anti-mouse Alexa 594. NPCs were analyzed for protein expression using the Protein Simple Jess automated western system per manufacturer protocol.

For YO-PRO-1 imaging, NPCs were washed once with HBSS –Ca –Mg (Gibco, catalog #14-175-095) with 20mM HEPES (Gibco, catalog #15-630-080) and preloaded with 10 μM YO-PRO-1 for 10 min. Imaging was performed on an Incucyte at 1 frame/min for 60 min total; after a 10-min baseline, 500 μM BzATP (Tocris, catalog #33-121) or vehicle was added. Per-cell uptake was extracted from segmented, drift-corrected movies using a custom Python pipeline (See Smith Github for code). Well-level fluorescence change was defined as post-BzATP response minus pre-injection baseline averaged over live cells and compared by Welch's t test.
